# Intercellular epigenomic signaling during B cell maturation

**DOI:** 10.1101/2024.05.24.595740

**Authors:** Kevin Ho Wai Yim, Alaa Al Hrout, Richard Chahwan

## Abstract

B cell maturation is crucial for effective adaptive immunity. It requires a complex signaling network to mediate antibody diversification through mutagenesis. B cells also rely on queues from other cells within the germinal center. Recently, a novel class of intercellular signals mediated by extracellular vesicles (EVs) has emerged. Studies have shown B cell EV-mediated signaling is involved in immune response regulation and tumorigenesis. However, the mechanistic role of B cell EVs is not yet established. We herein study the biological properties and physiological function of B cell EVs during B cell maturation. We use emerging technologies to profile B cell EVs surface marker signatures at the single particle level, molecular cargo, and physiological roles in B cell maturation. EV ncRNA cargo, characterized by RNA-seq, identified an EV-mediated novel non-coding RNA regulatory network for B cell maturation. A previously uncharacterized micro-RNA (miR-5099) in combination with a set of long non-coding RNA carried within B cell EVs is shown to be important for antibody diversification. The physiological role of EVs in B cell maturation is investigated using EV blockade assays and complementation studies using diverse EV sources further confirmed the physiological role and mode of action of EVs in B cell maturation.

## INTRODUCTION

Adaptive immunity is one of the main strategies used by vertebrates to safeguard from infections and attracted intensive research focus, especially with B cells, the factory of multiplex and high affinity antibodies to mediate specialized immune responses(1). The key processes involved in B cell antibodies diversification and affinity maturation are known as class switch recombination (CSR) and somatic hypermutation(2). CSR is a B cell specific intrachromosomal rearrangement of the switch region of the immunoglobulin (Ig) locus, from IgM to difference isotypes, which results in the generation of repertoire of antibodies (i.e., IgG1, IgG3, IgA, IgE) with different effector functions against invading pathogens. Defects in CSR in human leads to immunodeficiencies phenotypes such as hyper-IgM syndrome(3). The mechanism of CSR is a rather complex process that orchestrated by number of molecules and signaling factors. Murine primary splenic B cells have been widely used to study CSR regulation *ex vivo* with maximum CSR to IgG1 to around 20%, until the establishment of CH12F3, a high CSR efficiency (up to 60% CSR to IgA) cell line developed in 1996(2). However, the rationale for such significant difference in CSR efficiency between the two cell types is unclear. In addition, endogenous expression of CSR and DNA damage repair (DDR) associated genes between the two cell types cannot account for such differences.

Studies have shown non-coding RNAs (ncRNAs) are able to modulate B cell development and CSR(4). Small ncRNAs like miR-181b and miR-361 can silence AID expression in B cells and reduce CSR efficiency. Depletion of miR-155 in activated B cells reduces the production of IgG1+ plasma cells and memory B cells and therefore loss of IgG1+ antibodies(5). Extracellular vesicles (EVs), a new class of intercellular communication player, have been shown to deliver RNA and proteins, especially ncRNAs, from one cell and another to modulate gene expression in recipient cells(6). Although recent studies have shown EVs released by B cells can trigger primed CD4^+^ T cells proliferation and T_H_2 like cytokine production via delivery of MHC complexes(7, 8), inter B cell communication via EVs in physiological conditions requires further investigation. In this study, we attempted to understand the biological properties of B cell EVs during CSR that might contribute to physiological B cell development, and more importantly to determine whether the unknown difference in CSR efficiency between primary B cells and CH12F3 could be explained via EVs mediated signaling. Our findings, for the first time, showed significant increase of CSR of primary B cells via culturing in fresh CH12F3 derived conditioned medium, but not in freeze thawed medium which intact EVs were ruptured while other cell free proteins, nucleic acids, and cytokine factors were retained. Strikingly, the vice versa combination led to drastic reduction of CSR of CH12F3, suggesting the existence of EVs mediated CSR regulation in B cells. Differential expression of surface markers and EVs ncRNA cargo of B cell EVs during CSR was revealed with a specialized nanoparticle flow cytometer (NanoFCM®) and ncRNA-seq. Moreover, our data suggested effective EVs mediated CSR regulation requires the presence of EVs surface proteins and molecular cargo. Mechanistically, we identified a novel CSR suppressor, miR-5099, an uncharacterized miRNA that was only upregulated in stimulated B cell EVs, but not CH12F3 EVs. We also identified a long ncRNA CSR activator, Gm26917 that was upregulated in stimulated CH12F3 EVs. Taken together, these findings not only decipher the intrinsic difference in CSR efficiency between primary B cells and CH12F3, but also provide a novel perspective to study T cell independent CSR regulation via EVs.

## MATERIALS AND METHODS

### Cell cultures and *in vitro/ex vivo* CSR

CH12F3 cells were maintained and CSR assays performed as described previously(9). In brief, cells were stimulated with 1 ng/mL of recombinant human TGF-β1 (R&D Systems), 10 ng/mL of recombinant mouse IL-4 (R&D Systems) and 2 μg/mL of functional-grade purified anti-mouse CD40 (eBiosciences) and then analyzed by flow cytometry as described below. Splenic B cells were obtained from WT 8-wk-old female mice. Purified B cells were plated at a concentration of 0.5 × 10^6^ cells/ml and stimulated with either 50 µg/ml LPS (Sigma-Aldrich) or 50 µg/ml LPS and 50 ng/ml rIL-4 (R&D Systems) for 4 days. Animal protocols were reviewed and approved by the veterinary office of the canton of Zurich, Switzerland (213/2020).

### Isolation of EVs from cell cultures

Splenic B cells and CH12F3 were cultured for 2 days in RPMI 1640 plus particulate-depleted 10% FCS. FCS EVs were depleted using ultrafiltration filter (Amicon Ultra-15 100kDA) at 3,000 x g for 55 min. Culture supernatants were first spun at 400 × g for 5 min to deplete cells and then at 10,000 × g for 20 min to deplete residual cellular debris. EVs were isolated by 100,000 × g ultracentrifugation for 2 h at 4°C, and finally resuspended in PBS for further nanoflow analysis and functional assays. To deplete surface proteins on EVs surface, purified EVs in PBS were treated with Proteinase K (Thermo Fisher, 50 ng/µl) for 30 min at 37°C then quenched by incubation at 75°C for 5 min. To deplete EVs in suspension, purified EVs in PBS were ruptured by at least 3 cycles of freeze thaw or incubate with 1% Triton-X for 15 mins at room temperature.

### Transmission electron microscopy of purified EVs

To visualize EVs samples under the transmission electron microscope samples were transferred onto pioloform-coated EM copper grids by floating the grids on a droplet containing freshly prepared exosome placed on parafilm. After 5 min of incubation the grids were washed 3 x 5 min on droplets of deionized water before contrasting of bound exosomes in a mixture of 2% w/v methyl cellulose and 2% w/v uranyl acetate (mix 9:1) on ice for 10 min. Contrasted grids were then air dried in a wire loop before analysis using a JEOL JEM 1400 transmission electron microscope operated at 120kV. Images were taken with a digital camera (ES1000W, Gatan, Abingdon, UK). For immunogold labelling, exosomes were transferred onto EM grids as described above. After 3 x 5 min washes in deionized water unspecific binding sites were blocked with 0.5% fish skin gelatin (FSG) in PBS for 10 min before incubation with gold conjugated mouse IgM antibody (G5652, Sigma Aldrich) for 30 min. After 3 x 5 min washes in PBS the grids were incubated with 10nm protein A gold (BBI Solutions, Cardiff, UK) for 20 min before washing the grids again in PBS (6 x 5 min) followed by 10 x 1 min washes in deionized water. Exosomes were then contrasted and imaged as described above.

### Immunoblotting

Purified EVs were lysed with RIPA buffer and heated at 95°C for 10 min before being loaded for electrophoresis. The membrane was incubated with a primary rabbit anti-CD63 antibody (10628D, Invitrogen), anti-Alix (Ma1-83977, Invitrogen), anti-CD19 (14-0194-82, Invitrogen), anti-AID (14-5959-80, Invitrogen) and anti-Golgin 97 (PA5-30048, Invitrogen) overnight at 4°C on a shaker. Membranes were washed with TBST three times at 15-min interval. The goat anti-rabbit IRDye 680 RD secondary antibody (1:1,000 dilution, Li-COR, catalogue number: 925–68071) was then incubated with the membrane for 1 h at room temperature. Blots were imaged using the Li-COR Odyssey scanner after wash.

### Immunostaining and flow analysis of cells

Surface IgM, IgA, IgG1 on CH12F3 and primary B cells were determined by BD FACSCanto II cell analyzer with APC-eFlour 780 anti-mouse IgM (47-5790-82 eBioscience), FITC anti-mouse IgA (11-4204-81, eBioscience) and PE anti-mouse IgG1 (12-4015-82, eBioscience) antibodies. Cells were washed in PBS at 300 x g for 5 min. Antibody staining was performed at the concentration of 1 in 200. Stained cells were washed once with PBS before acquisition. Gating was based on single cells followed by cell viability staining (65-0866-14, Thermo Fisher). All data analysis of cells and EVs were carried out with FlowJo (TreeStar) software and Graphpad Prism 9 software.

### Immunostaining and nanoflow analysis of purified EVs

Purified EVs were stained with FITC anti-mouse IgA (11-4204-81, eBioscience), FITC anti-mouse IgG1 (11-4015-82, eBioscience), PE anti-mouse IgM (12-5790-82, eBioscience), FITC Rat IgG2b, κ Isotype Ctrl (400633, Biolegend), FITC anti-mouse CD9 (124807, Biolegend), FITC anti-mouse CD19 (302206, Biolegend) and followed by 100,000 x g for 2 h at 4°C to remove unbound antibodies. Stained EVs were resuspended in 100 µl of 0.22 µm filtered PBS and subjected to flow analysis by NanoFCM Flow Nanoanalyzer. Monodisperse silica nanoparticles of four assorted sizes, with modal peak sizes of 66 nm, 91 nm, 113 nm and 155 nm were used as the size reference standard to calibrate the size distribution of EVs. Unless stated otherwise, all samples were recorded in 1 minute with the range of 2,000 to 12,000 events per minute to minimize the swarm detection effect since there is enough time/space between each particle measurement as evinced by the event burst trace which shows separate measurements of particles. The steady flow of the system allows for comparison of particle detection rate to a concentration standard (a stable 250nm silica bead of 1.87e10/ml (batch variable)). The particle concentration is then calculated, including the dilution factor, in particles/ml. All data analysis of cells and EVs were carried out with FlowJo (TreeStar) software and Graphpad Prism 9 software.

### RNA extraction

Cell and EV total RNA were extracted by Trizol^®^ (Invitrogen) and purified by RNeasy^®^ Mini Kit (Qiagen). Briefly, cell or EV pellets 700 µl Trizol^®^. Subsequently, 140 µl of chloroform was used for phase separation. Trizol^®^ and chloroform mix were centrifuged at 12,000 x g for 15 mins for phase separation. 300 µl RNA-containing top clear phase were transferred to a fresh tube and 400 µl absolute ethanol was added and mixed. 700 µl of sample were transferred to RNeasy spin column. DNAse treatment was performed after first 350 µl RW1 washing step. Finally, RNA was eluted in 30 µl RNase-free water. RNA concentration was assessed using a NanoDrop 2000 spectrophotometer (Thermo Scientific, Waltham, MA, USA). The RNA yield and size distribution were analyzed using an Agilent 2200 Tapestation with high sensitivity RNA Screentape (Agilent Technologies, Foster City, CA, USA).

### RNA-seq library preparation, next-generation sequencing and data processing

For small RNA library preparation, RNA aliquots were used for library preparation using NEBNext Multiplex Small RNA library preparation kit (New England Biolabs, Ipswich, MA, USA). The PCR amplified cDNA construct (from 140–160 bp) was purified using a QIAquick PCR Purification kit (Qiagen). The purified cDNA was directly sequenced using an Illumina MiSeq 2000 platform (Illumina, San Diego, CA, USA).

For long non-coding RNA library preparation, libraries were constructed using Ribo-Zero Magnetic Gold Kit (Human) (Illumina, San Diego, CA, USA) and NEBNext® Ultra™ RNA Library Prep Kit for Illumina (New England Biolabs) according to the manufacturer’s instructions. Libraries were tested for quality and quantified using qPCR (Kapa Biosystems, Woburn, MA, USA). The resulting libraries were sequenced on a HiSeq 2500 instrument that generated with either single -end (CH12F3) and paired-end (primary B cells) reads of 100 nucleotides.

Raw sequencing reads were checked for potential sequencing issues and contaminants using FastQC (https://www.bioinformatics.babraham.ac.uk/projects/fastqc). Adapter sequences, primers, number of fuzzy bases (Ns), and reads with quality scores below 30 were trimmed using fastq-mcf. Clean reads were aligned to the mouse genome (GRCm38/mm10) using the TopHat 2.0 program, and the resulting alignment files were reconstructed with Cufflinks (10). The transcriptome of each sample was assembled separately using Cufflinks 2.0 program. All transcriptomes were pooled and merged to generate a final transcriptome using Cuffmerge. After the final transcriptome was produced, Cuffdiff was used to estimate the abundance of all transcripts based on the final transcriptome to generate read counts for each sample.

### Gene expression quantification by RT-qPCR

800 ng total RNA was converted into cDNA using High-Capacity cDNA Reverse Transcription Kit according to manufacturer’s protocol. qPCR was performed using HOT FIREPol^®^ EvaGreen^®^ qPCR Mix Plus no ROX (Solis Biodyne) and analyzed with a Bio-Rad CFX384 Touch Real-Time PCR Detection System.

### Sequencing data analyses and statistical methods

Read counts of each sample were subjected to cluster analysis and differential expression (DE) analysis using RNA-seq 2G (11). |fold-change| ≥1, P value ≤0.05 and false discovery rate (FDR) ≤0.05 was considered statistically significant DE ncRNAs. DE ncRNAs expression in different immune cell types was determined using My Geneset ImmGen(12). Interaction and gene targets of identified DE ncRNAs in cells and paired EVs were predicted by miRNet and ENCORI(13, 14). Motif analysis were performed using XSTREME motif discovery and enrichment analysis software(15). Briefly, FASTA sequence of miRNAs in each sample (e.g. cells v.s. paired EVs; unswitched EVs v.s. switched EVs) were used as input for RNA motifs discovery and enrichment analysis (Ray2013 All Species), Analytical parameters were set as following, E-value ≤ 10, motif width between 4 and 15, and alignment starting from left ends. Motifs with over 50% true-positive and 0% false-positive enrichment were selected as unique motifs for indicated sample.

### EVs blockade and gene silencing/overexpression assay

10 µM and 5 µM GW4869 was used to block EVs production in CH12F3 and primary B cells respectively prior to cytokine activation. Expression of target ncRNAs in CH12F3 and primary B cells were inhibited by 2 µM miRCURY LNA miRNA Inhibitors (339131, Qiagen), anti-miR-5099 (GGAGCACCACATCGATCTAA-FAM); antimir-control (TAACACGTCTATACGCCCA-FAM), Antisense LNA GapmeR for Gm26917 was designed using the LNA GapmeR designer (Qiagen). Transfection efficiency was quantified by FAM signal intensity increase prior to cytokine activation. Overexpression of miR-5099 in CH12F3 was achieved by transfecting 2 µM of miR-5099 miRCURY LNA miRNA mimic (Qiagen) by electroporation using Lonza 4D-Nucleofector and cell line SF kit following manufacturer’s protocol.

## RESULTS

### Extracellular constituents are required for high CSR efficiency in B cells

Primary splenic B cells has been the classical study model for CSR, until the emergence of CH12F3, a high switching frequency and single isotype committed (IgA exclusively) cell line established by Honjo et. al(2). We compared the CSR efficiency of primary B cells (from IgM^+^ to IgG1^+^) and CH12F3 (from IgM^+^ to IgA^+^) upon cytokine stimulation in our hands to the data from other publications (Fig. 1A), a significant difference in switching efficiency was observed, with ∼20% switching in primary B cells and ∼60% switching in CH12F3. However, the intracellular molecules responsible for such differences has not been identified so far. Therefore, we explored the possibilities of extracellular mediated CSR regulation by culturing primary B cells in conditioned medium (CM) from stimulated CH12F3, and vice versa (Fig. 1B). CM from both cell types were either used directly or after three freeze thawing cycles to retain only potential functional cytokines, proteins, and nucleic acids, but not intact extracellular vesicles (EVs) (Fig. S1)(16–20). Strikingly, for the first time, significant increase in primary B cells CSR to IgG1^+^ was observed in CH12F3 CM condition, but not in freeze thawed CM, compared to autologous CM control (Fig. 1C). Moreover, CH12F3 CSR to IgA^+^ was drastically decreased only in fresh primary B cell CM condition compared to autologous CM control (Fig. 1D). Loss of function in freeze thawed CM highlights EVs contribution and neither free floating proteins nor nucleic acids retained in freeze thawed CM were able to provide comparable regulatory effects. These novel findings were showing the existence of extracellular mediated CSR regulation in B cells independent of T cell signaling in physiological settings for the first time.

**Figure 1.**
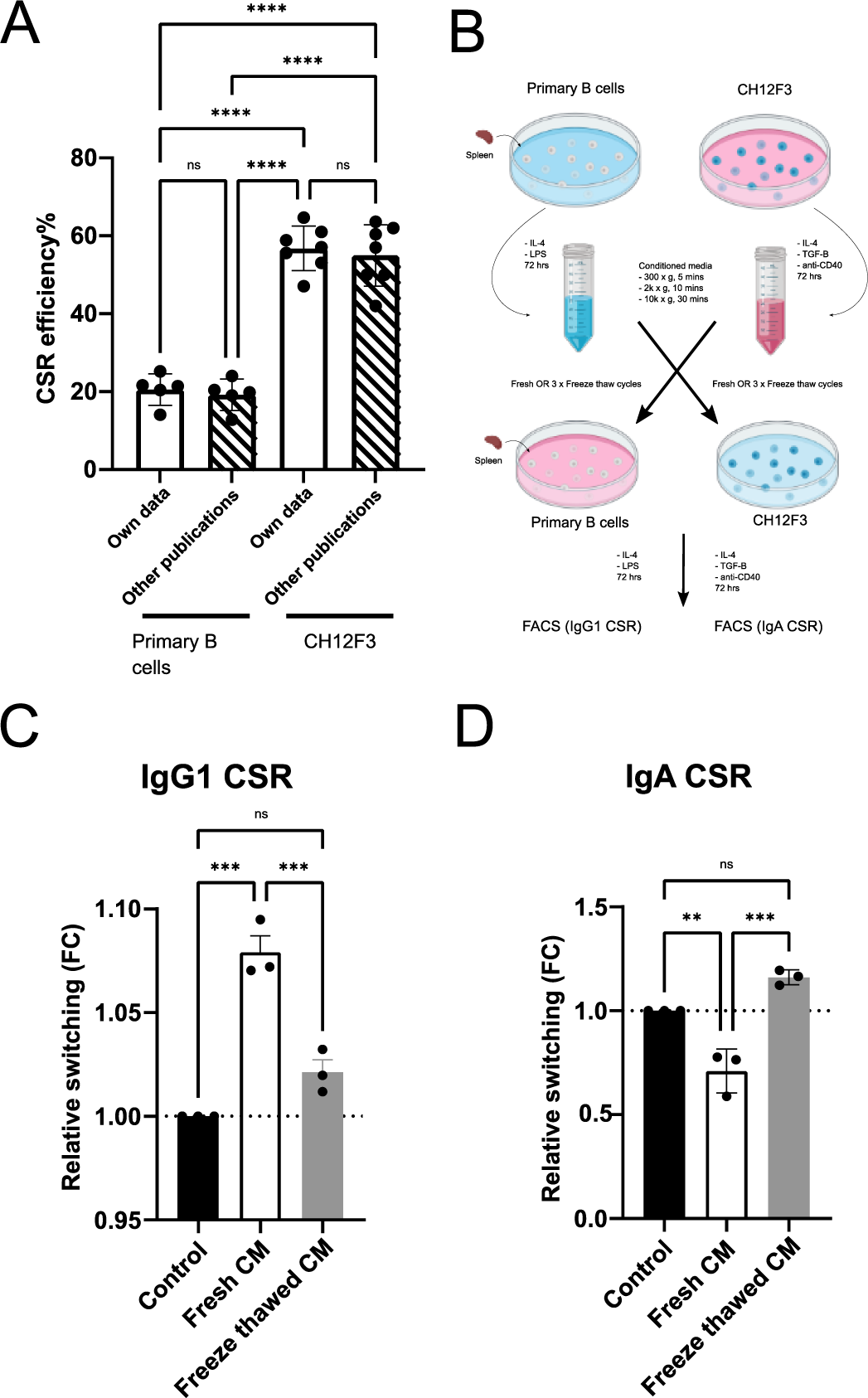
Extracellular constituents from different B cell lines regulate CSR. (A) Comparison of CSR efficiency in primary splenic B cells and CH12F3 from own data and other published data. (B) Schematic outline of conditioned medium exchange experiment. (C & D) IgG1 CSR of primary splenic B cells (C) and IgA CSR of CH12F3 (D) in the presence of denoted conditions. Representative of three experiments, mean ±SD shown in columns by one-way ANOVA with Tukey’s multiple comparison test (*p<0.05; **p < 0.01; ***p < 0.001).

### CSR causes differential surface proteins and cargo profiles on released EVs

To better understand how CSR affects the biological properties of B cell derived EVs, we studied B cell EVs derived from B cells with or without cytokine stimulation. Since culturing primary B cells *ex vivo* requires stimulation for survival after purification from spleens, thus, it was not possible to isolate EVs from unstimulated primary B cells per se. For this matter, we used CH12F3 EVs as our study model here since they undergo CSR in higher efficiency compared to other cell lines, and more vitally, the possibility to isolate EVs from unstimulated state. EVs from unstimulated and stimulated CH12F3 were prepared as described (Fig. 2A). In addition, fetal calf serum used in cell cultures contained significant amount of heterologous EVs (hEVs) that, we showed here for the first time, could affect B cell survival and from reaching their maximum CSR efficiency (Fig. S2). Western Blot analysis of CH12F3 and released EVs with or without stimulation showed a differential protein expression profile. Interestingly, for the first time we observed that, the pan B cell marker, CD19 was more expressed in EVs compared to cell, suggesting an active cargo shuttling into secreted EVs. The classical EV tetraspanins marker, CD63 was not highly expressed in CH12F3 EVs which correspond to other publication(21). One of the most vital mediators for CSR, AID protein, was not or lowly detectable in stimulated CH12F3 EVs, suggesting the CSR regulatory function by EV signaling was not mediated by direct delivery of AID (Fig. 2B). Transmission electron microscopy (TEM) analysis in our data showed, for the first time in the field, pockets of exosome-sized vesicles being secreted by B cells. Using the MAPS software (Thermo Scientific™) to perform high resolution analysis, the sizes and quantification of individual vesicles as well as the entire pockets were determined. Strikingly, in one TEM cross section (70 nm), vesicles pockets were found in 20 out of 158 cells which we estimated around 10 pockets being secreted by an actual B cell (between 7-10 µm) at any time point. Furthermore, the presence of B cell derived EVs was readily detected in mice serum using anti-mouse IgM immunogold labeling, suggesting their biological relevance in physiological conditions (Fig. 2C). Next, single nanoparticle flow analyzer, NanoFCM® (nFCM) was being utilized the first time in the B cell EV field, to profile surface markers, as well as sizes and quantity of CH12F3 EVs with or without stimulation. Using known size silica beads (r index: 1.46) as reference, the mean estimated sizes of unstimulated and stimulated CH12F3 EVs were ranging between 80 nm to 90 nm, which was comparable to our TEM data (Fig. 2D, Fig. S3). The ratio between number of CH12F3 and their EVs were quantified, align with previous studies(7, 22), stimulated CH12F3 released 10-fold more EVs compared to unstimulated CH12F3, suggesting a correlation between cellular stress and EVs secretion in B cells (Fig. 2E). Interestingly, ratio between total protein and number of EVs in stimulated CH12F3 EVs was around 10-fold less compared to unstimulated CH12F3 EVs, indicating activated B cells selectively shuttle only the critical functional protein cargo into secreted EVs in exchange of higher EVs secretion. (Fig. 2F). Surface marker profile of EVs derived from CH12F3 with or without stimulation was determined, human embryonic kidney 293A (HEK293A) EVs were used as negative staining control. Potential fluorophores spillovers (co-labeling of FITC and PE conjugated antibodies) were also assessed using single stained EVs (Fig. S4). Interestingly, only 10% of CH12F3 EVs express classical tetraspanins marker, CD9, and half of which also express IgM. CD9 expression on EVs was not significantly altered during CSR. The low expression of CD9 in the total CH12F3 EV population suggested that tetraspanins might not always be a generic marker of EVs purification or capture for FACS, especially for B cell EVs, and therefore, a more expressed marker will be needed for B cell EV studies. Pan-B cell marker, CD19 were expressed on 35% of total EVs population from stimulated CH12F3, with 30% only CD19^+^ and 5% IgM^+^CD19^+^. Here, we observed for the first time, a 3-fold significant increase in CD19 expression on EVs during CSR, moreover, such increment was not detectable by Western Blot analysis. Surface IgM and IgA on EVs were compared during CSR. Amount of IgM^+^ EVs were consistent during CSR, however, IgA^+^ EVs were 5-fold more in CH12F3 after stimulation (Fig. 2G). Similar results were also observed between unswitched (24 hours post stimulation) and switched (72 hours post stimulation) primary B cells derived EVs, including significant increased total EVs secretion, elevated expression of CD9, CD19, and CSR specific IgG1 on secreted EVs during CSR (Fig. S5) Taken together, these findings provided a novel comprehensive foundation for the field to better understand physiological EVs mediated signaling in B cells development.

**Figure 2.**
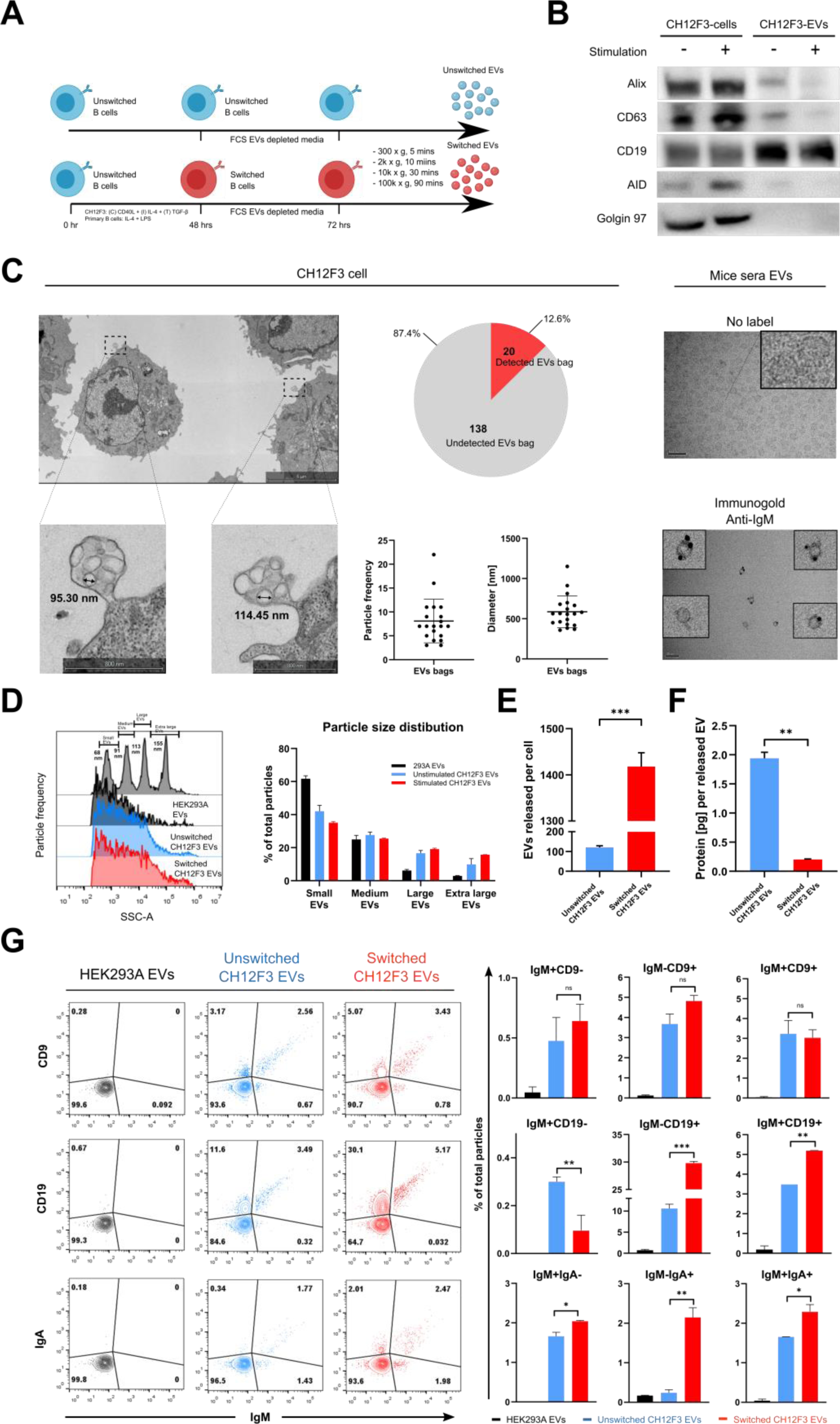
Multiplex characterization of B cell EVs during CSR. (A) Schematic outline of cell stimulation and EVs collection. (B) Western blot analysis of denoted protein markers from CH12F3 and derived EVs during CSR. (C) Left: transmission electron microscopy analysis of CH12F3, with suspected EVs bags morphology and quantification. Right: Immuno-gold labeling of IgM in mice serum EVs. (D) Representative side scatter plot and estimated size distribution quantification of CH12F3 EVs during CSR, using silica sizing beads with mix of 66 nm (small), 91 nm (medium), 113 nm (large), and 155 nm (extra-large) (r index = 1.46). (E) Quantification of estimated EVs particles released per cell via dividing the total number of particles by number of parental cells. (F) Quantification of estimated EVs proteins per released particles via dividing the total amount of EVs proteins by number of particles. (G) Representative FACS plots and quantification of denoted markers of CH12F3 EVs during CSR. Representative of three experiments, mean ±SD shown in columns by one-way ANOVA with Tukey’s multiple comparison test (*p<0.05; **p < 0.01; ***p < 0.001).

### Differential EVs ncRNAs repertoire in primary B and CH12F3 EVs

Increasing number of evidence has demonstrated different cell types actively and selectively shuttle ncRNAs cargo into secreted EVs to perform intercellular signaling in recipient cells(6). To explore the possibility of EV ncRNAs mediated regulation in B cell CSR, we performed RNA-seq analysis. Total RNA from the cells and EVs were extracted from unswitched and switched primary B cells and CH12F3 for exploratory screening (Fig. 3A). Small RNA and lncRNA libraries were first prepared from extracted total RNA and subjected to RNA-seq analysis. The abundance of miRNAs and lncRNAs in CH12F3 and primary B cell derived EVs with over 20 reads were quantified. LncRNAs were more represented in CH12F3 EVs compared to miRNAs in unswitched state (243 lncRNAs vs 51 miRNAs), and further enriched after stimulation of parental cells (525 lncRNAs vs 55 miRNAs). However, in primary B cell EVs, number of miRNAs and lncRNAs were similar in unswitched state (45 lncRNAs vs 49 miRNAs), and miRNAs were more enriched after switching (36 lncRNAs vs 106 miRNAs) (Fig. 3B). Apart from ncRNAs abundance, ncRNAs expression between primary B cell and CH12F3 EVs was also significantly different as shown in the log_2_ transformed heatmap (Fig. 3C), suggesting the two cell types have a vastly different EVs cargo packaging regimen and thereby exert differential effect in CSR regulation.

**Figure 3.**
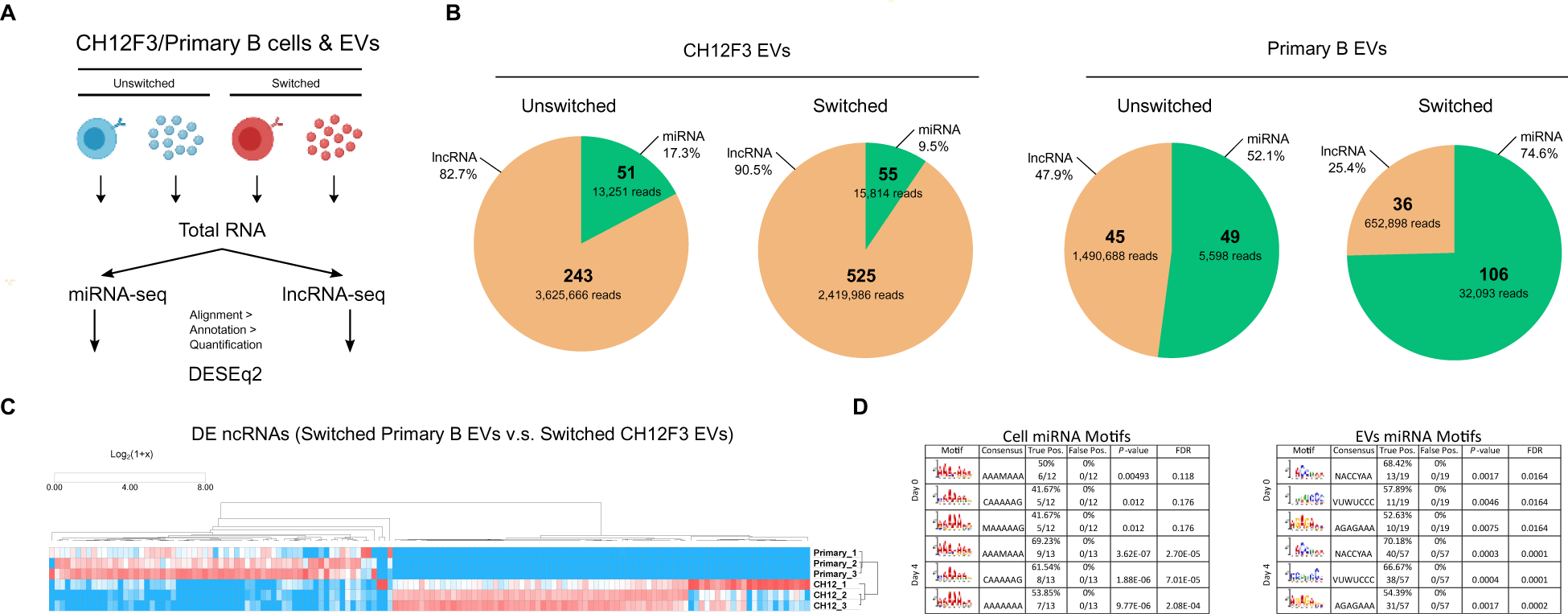
Varied non-coding RNA profile in primary B cells EVs and CH12F3 EVs during CSR. (A) Schematic outline of RNA-seq analysis of molecular cargo in EVs and paired cells. (B) Quantification of non-coding RNA (ncRNA) reads (>30 reads) from EVs in denoted conditions. (C) Expression heatmap of differentially expressed ncRNAs between primary B EVs and CH12F3 EVs. Hierarchical clustering by one minus spearman rank correlation of log_2_(1+x) transformed read counts. (D) Shared motifs between primary B cells and CH12F3 over-represented in miRNAs preferentially sorted into EVs or retained in cells in indicated days post-CSR. Table showing the cell retaining (left) and EVs sorting (right) motifs identified in both primary B cells and CH12F3. For each motif, the percentage of true positive (primary sequences matching indicated motif), false positive (control sequences matching indicated motif), P-value, false discovery rate (FDR are displayed.

To understand if B cells also selectively package miRNAs into secreted EVs as reported in recent study (23), differentially expressed miRNAs in paired cells and EVs, pre- and post-CSR from both CH12F3 and primary B cells were subjected to Simple Enrichment Analysis (15) to identify motif enrichment in EVs compared to parental cells (Fig. S6). Unique motifs enriched in cells or paired EVs were first selected and those expressed in at least 50% of input miRNAs and p-value < 0.001 were then included in the list of motif candidates. Finally, for cells or EVs enriched motifs, only the ones that present in both primary B cells and CH12F3 throughout CSR were considered as potential conserved cell retaining and EVs sorting motifs in B cells. In cell miRNA motifs, polyA sequence motifs were highly overrepresented throughout CSR in both primary B cells and CH12F3 and even more significant four days post CSR. Interestingly, three unique EVs enriched motif consensuses, NACCYAA, VUWUCCC, and AGAGAAA were identified throughout CSR in both EVs sources, suggesting the existence of sequence motif-based miRNAs sorting mechanism in B cells (Fig. 3D).

### EVs mediated CSR regulation is surface and cargo dependent

To determine the mode of EVs mediated CSR regulation, EVs production in CH12F3 and primary B cells was first inhibited by GW4869(24–26), a ceramide synthesis inhibitor prior to cytokine stimulation for CSR. At the same time, to validate the contribution of EVs in such regulation, but not other exogenous factors (i.e. proteins or cytokines), complement of autologous EVs in four forms, fresh, proteinase-K treated (depletion of protein aggregates and EVs surface proteins)(27), triton-X-treated, and freeze-thawed (depletion of intact EVs) (Fig. 4A, Fig. S4) was performed. As expected, depletion of released EVs in CH12F3 resulted in reduction in CSR to IgA, however, CSR to IgG1 in primary B cells was enhanced compared to DMSO control. It further proved the existence of differential regulation mediated by EVs from the two cell types. Furthermore, complement of fresh autologous EVs was able to rescue the phenotype observed above, however, proteinase-K treated EVs only showed half rescue compared to fresh EVs, and triton-X treated EVs were not able to rescue the CSR reduction. Taken together, CH12F3 EVs are promoting CSR while primary B cell EVs are suppressive for CSR, and such EV mediated CSR regulation is likely to be a dual step process that relies on EVs surface proteins, possibly for docking and signal triggering, as well as molecular cargo delivered by intact EVs for gene modulation in recipient cells (Fig. 4B, 4C).

**Figure 4.**
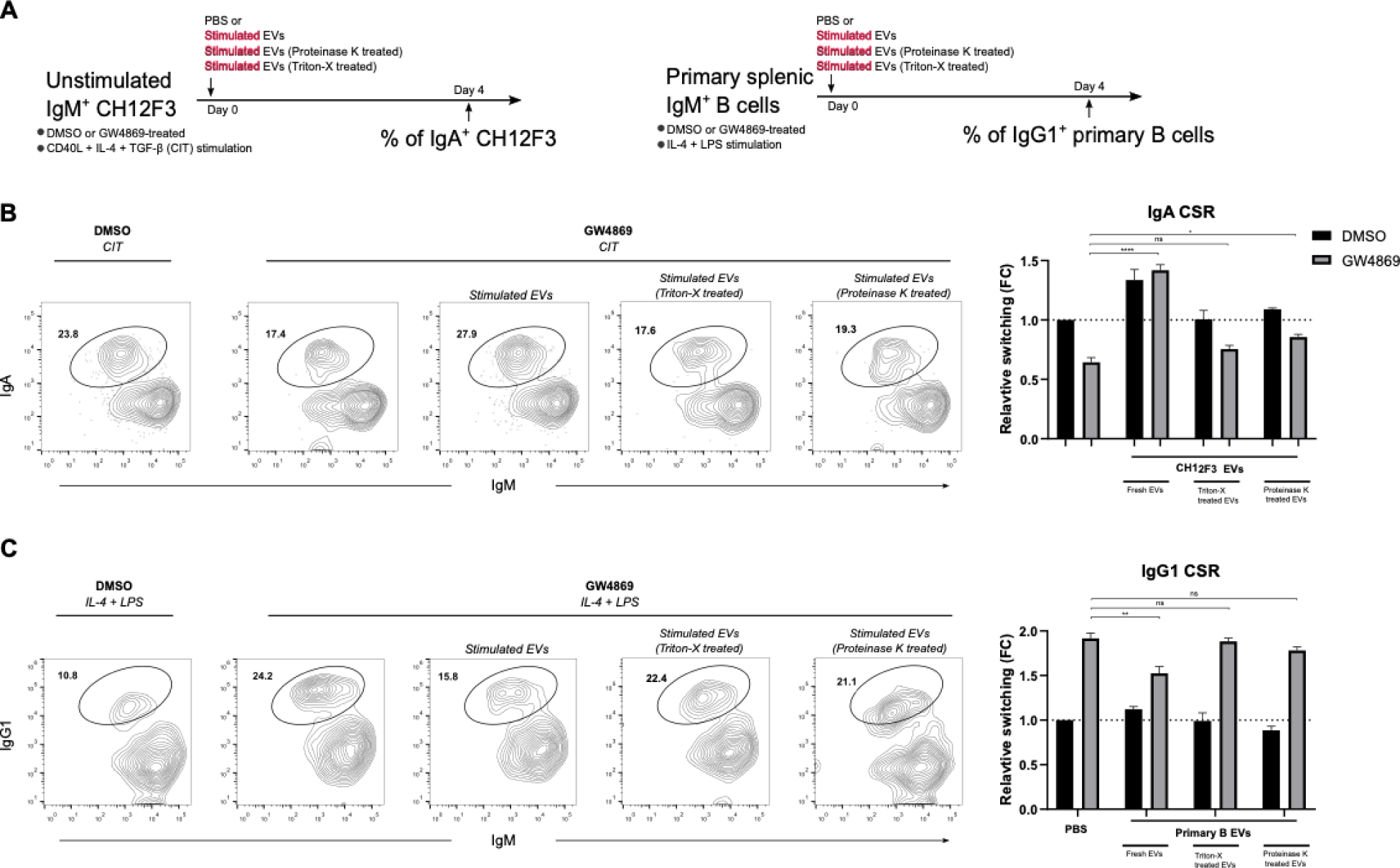
Varied CSR regulatory effect arise from different source of EVs. (A) Schematic outline of functional phenotypic assay of EVs mediated CSR in CH12F3 (left) and primary B cells (right). (B and C) Representative FACS plots and quantifications of CSR efficiency in either DMSO (mock), GW4869 (EVs release inhibitor) with the addition of fresh EVs, proteinase-K treated EVs (surface antigens shaved), freeze-thawed, and Triton-X treated EVs (ruptured) in CH12F3 and primary B cells. Representative of three experiments, mean ±SD shown in columns by two-way ANOVA with Tukey’s multiple comparison test (*p<0.05; **p < 0.01; ***p < 0.001).

### EVs ncRNA cargo are responsible for CSR regulation

EVs ncRNAs cargo have been shown to modulate immune responses *in vitro* and *in vivo*(6). To identify the responsible EV ncRNAs in CSR modulation observed in figure 1c and 1d, we focused on the significant DE miRNAs and DE lncRNAs in unswitched and switched parental cells and secreted EVs (Fig. 5A). miR-5099, a miRNA enriched in germinal center B cells compartment relative to known high expressers, AID and low expressers, Rag2 (Fig. S7) (according the Immunological Genome (ImmGen) consortium studies)(12). miR-5099 has no reported function in B cells nor other biological processes, were highly upregulated (Log_2_ fold change = 3.653) in primary B EVs after stimulation. Interestingly, miR-5099 was downregulated (Log_2_ fold change = −5.223) in the parental cells at the same time after stimulation (Fig. 5B), suggesting primary B cells actively package miR-5099 into released EVs. To identify potential linkage to CSR associated genes, we employed ENCORI database to explore the interactive targets of miR-5099 (table. S1)(14). Surprisingly, one of the two lncRNAs is upregulated in switched CH12F3 and secreted EVs, Gm26917 had the highest number of interactions and lowest minimum free energy of such RNA-RNA pairs. To probe the functional role of these two ncRNAs in CSR regulation, CH12F3 were treated with anti-miR-5099, miR-5099 mimic, and antisense LNA GapmeR for Gm26917 prior to CSR. Both anti-miR-5099 and GapmeR Gm26917 were labeled with FAM that enable us to directly quantify the corresponding uptake in cells by FACS. miR-5099 mimic was co-transfected with GFP plasmid by nucleofection to confirm transfection efficiency (Fig. S8). Strikingly, inhibition of miR-5099 leads to significant increase in both cell types compared to control, suggesting miR-5099 is a pan CSR suppressor in B cells (Fig. 5C, S8). To confirm the CSR suppressing role of miR-5099, CH12F3 were transfected with miR-5099 mimic and subjected to CSR. Overexpression of miR-5099 led to drastic CSR reduction compared to control (Fig. 5D), this further suggested miR-5099 is used by primary B cells to actively regulate CSR via EVs mediated signaling (Fig. 6). Since Gm26917 is one of the most significant DE lncRNAs in CH12F3 EVs during CSR and being the top interactive target of the CSR suppressor miR-5099, we tested whether Gm26917 also involves in CSR regulation. Notably, silencing of Gm26917 by antisense LNA oligonucleotides in CH12F3 has reduced CSR by 40%, suggesting the CSR promoting role of Gm26917. Inhibition of mir-10b, one of the upregulated miRNAs upregulated in CH12 EVs post CSR did not affect CSR efficiency, showing the specificity of identified ncRNAs (Fig. S9). To confirm the interaction between miR-5099 and Gm26917 and associated CSR phenotypes, the expression of Gm26917 and AID was quantified in anti-miR-5099 treated, miR-5099 mimic, and GapmeR Gm26917 treated CH12F3 during CSR by RT-qPCR (Fig. 5F. Inhibition of miR-5099 reduced AID expression and increased Gm26917 expression, showing the CSR regulatory role of miR-5099 and the interaction between miR-5099 and Gm26917. In GapmeR Gm2617, the reduced Gm26917 expression confirmed the effective silencing of Gm26917. Reduction of AID in GapmeR Gm26917 treated CH12F3 suggested its CSR promoting role. In contrast, overexpression of miR-5099 reduced Gm26917 expression and increased AID expression (Fig. 5FG), suggesting that Gm26917 could potentially be a suppressor of the CSR suppressor, miR-5099.

**Figure 5.**
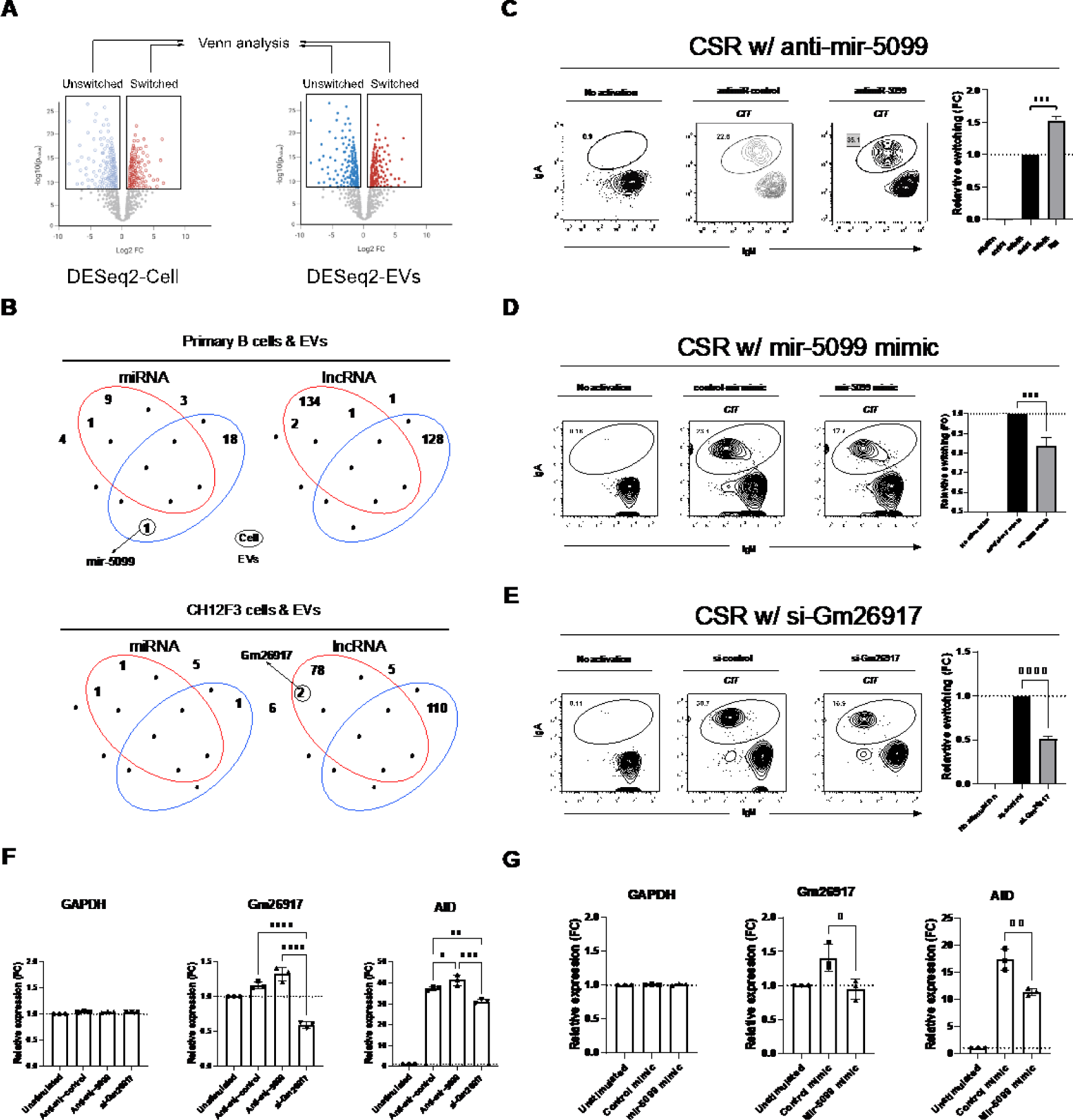
EVs regulate CSR via delivery of specific non-coding RNA. (A) Schematic outline of DESeq2 and Venn analysis of molecular cargo in EVs and paired cells during CSR. (B) Venn diagram analysis of differentially expressed ncRNA (Fold change ≥ 2, FDR < 0.05) in EVs and paired cells during CSR. (C-E) Representative FACS plots and quantifications of CSR efficiency in CH12F3 treated with anti-miR-5099 (C), miR-5099 mimic (D), and GapmeR Gm26917 (E) compared to respective negative controls. Representative of three experiments, mean ±SD shown in columns by one-way ANOVA with Tukey’s multiple comparison test (*p<0.05; **p < 0.01; ***p < 0.001). (F and G) Quantifications of relative expression of denoted genes in CH12F3 treated with anti-miR-5099 and GapmeR Gm26917 (F), and miR-5099 mimic compared to control (G), mean ±SD shown in columns by one-way ANOVA with Tukey’s multiple comparison test (*p<0.05; **p < 0.01; ***p < 0.001).

**Figure 6.**
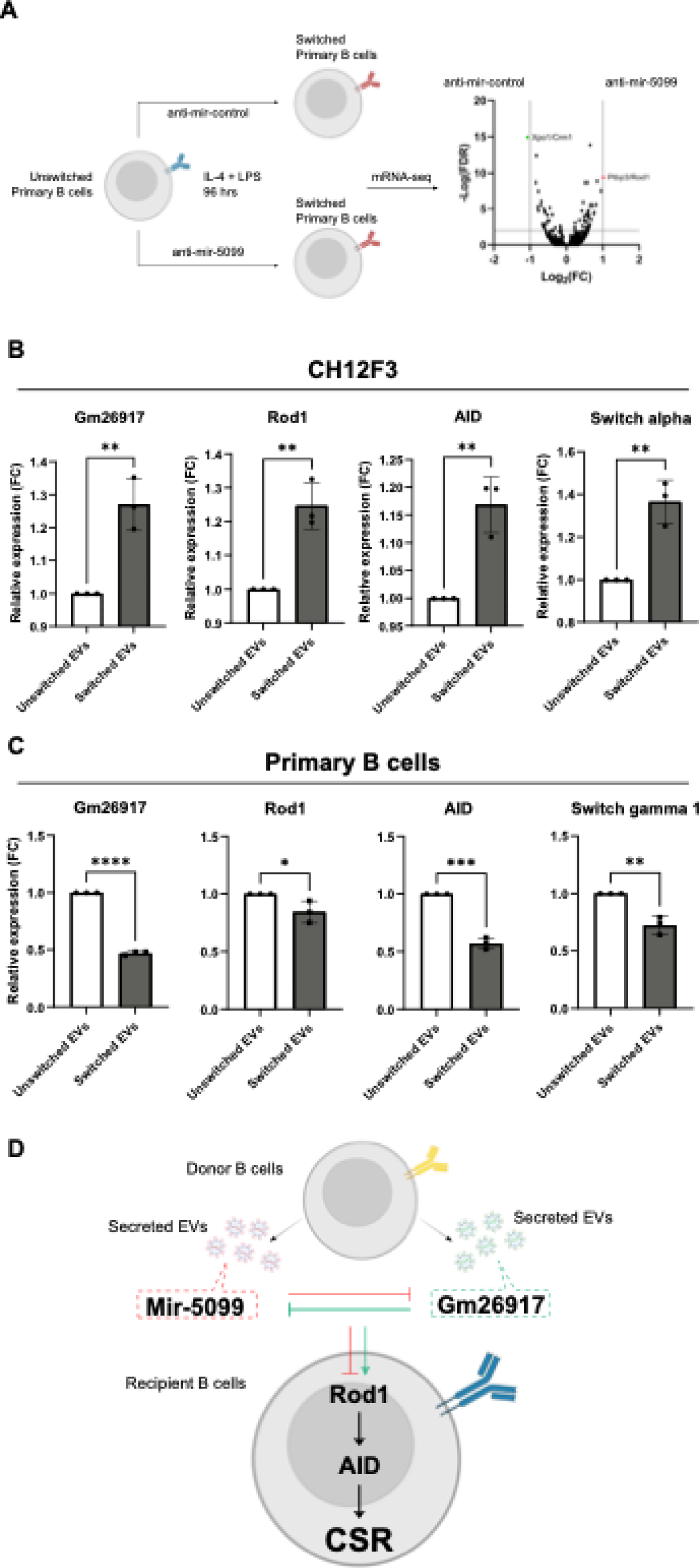
B EVs ncRNAs regulate CSR through PTBP3/Rod1 pathway. (A) Schematic outline of mRNA-seq of primary B cells treated with anti-miR-5099 and volcano plot showing differentially expressed downstream mRNAs in anti-miR-5099 treated cells compared to control. (B and C) Quantifications of relative expression of denoted genes in recipient CH12F3 (B) and primary B cells (C) in the presence of autologous EVs (unswitched and switched parental cells) 48 hours post stimulation. Mean ±SD shown in columns by one-way ANOVA with Tukey’s multiple comparison test (*p<0.05; **p < 0.01; ***p < 0.001). (D) Proposed mechanism of B EVs mediated CSR regulation.

### EV miR-5099 suppress CSR via interaction with ROD1

To understand the downstream mechanism of CSR suppressor miR-5099, RNA-seq was performed with stimulated primary B cells treated with either anti-mir-control and anti-miR-5099. Interestingly, transcript of the RNA binding protein, ROD1 (also known as PTBP3) was upregulated in anti-miR-5099 treated primary B cells compared to control (Fig. 6A). ROD1 has already been reported to interact with AID and serves as an essential guiding system for AID to the Ig loci during CSR in *ex vivo* studies(28). The other two members of this gene family PTBP1 and PTBP2 are also known to be essential for B cell maturation and development by controlling mRNA abundance and alternative splicing of vital cell cycle regulators(29–31). To confirm the interaction between miR-5099 and ROD1 at the transcription level, the expression of ROD1 was quantified in anti-miR-5099- and miR-5099 mimic-treated B cells after CSR. Upon treatment with anti-miR-5099, ROD1 expression was increased by 15% compared to control. Whilst in miR-5099 mimic-treated CH12F3, ROD1 expression was decreased by 25% compared to control. ROD1 expression was also decreased upon silencing of Gm26917, suggesting Gm26917 indirectly affect ROD1 expression possibly via interaction with miR-5099 (Fig. S10). Next, we attempted to confirm that this novel CSR regulation is mediated by EVs signaling, expression of Gm26917, ROD1, AID, switch alpha (CH12F3), and switch gamma 1 (primary B cells) were quantified in recipient CH12F3 and primary B cells in the presence of autologous EVs (either from unswitched and switched parental cells) 48 hours post stimulation. As expected, Gm26917, ROD1, AID, and switch alpha showed significant increase in CH12F3 with the presence of autologous EVs from switched parental cells compared to unswitched control (Fig. 6B). In contrast, expression of Gm26917, ROD1, AID, and switch gamma 1 were reduced in primary B cells with the presence of autologous EVs from switched parental cells compared to unswitched control (Fig. 6C). Overall, our findings have revealed a novel intercellular epigenomic signaling form of CSR regulation via EVs in physiological B cell maturation (Fig. 6D).

## DISCUSSION

CSR is a crucial process for diversification of antibodies effector functions in adaptive immunity. The two main study models, primary splenic B cells and CH12F3, representatives of healthy and oncogenic respectively, have a significant variance of CSR efficiency that has not be explained by endogenous proteomic, genomic, transcriptomic, nor epigenetic studies. Indeed, understanding of the molecular mechanism contributing to such variance would enhance our knowledge in CSR regulation and treating CSR defects associated diseases. From the conditioned medium exchange assay in figure 1, we revealed a novel EVs mediated intercellular epigenomic CSR regulatory network in physiological conditions independent of T cells signaling. These findings contributed to our understanding of the significant variance in CSR efficiency between primary splenic B cells and CH12F3, and provide novel insights in how B cells with different background (i.e., healthy v.s. cancerous) utilize EVs signaling to tightly regulate their autologous activation equilibrium. Our findings hold a promising potential of developing into an early diagnostic and alternative therapeutic applications in B cell associated autoimmune disorders and cancers.

B cell associated EVs studies in physiological conditions are still in a growing phase compared to those in other diseased models and the tools for studying EVs have evolved tremendously in recent years. Therefore, to gain more accurate knowledge in our studied EVs, we applied the cutting-edge single particle nanoflow based technology combining with conventional imaging techniques to establish the biological background of B cell derived EVs during physiological CSR. From our TEM analysis of CH12F3, we identified exosome-sized (within 40 to 150 nm) EVs being secreted in an encapsulated bag format by 12% of the 2D imaged cells in 100^th^ of an actual cell cross-section (70 nm). Considering the cells are 3D structured in reality, every cell is estimated to secrete 10 bags of EVs in any given time point, indicating the constitutive intercellular EVs communication between CH12F3. Such form of particle release was also observed in viral infected T cells, colorectal cancer cells, cardiac telocytes and gastrointestinal stromal tumors(32–35), we are the first to report such mode of secretion of EVs under physiological condition from B cells, possibly to enhance protection and stability for the individual EVs in the extracellular space. Since the mode of EVs biogenesis and secretion are not yet fully understood in B cells, our work displayed a clear visualization of such processes and laid foundation for future EVs studies focusing on B cell EVs biogenesis and secretion pathways.

Apart from morphological analysis, we attempted to determine the biological properties in B cell EVs during CSR using NanoFCM. Previous studies were mostly based on bulk EVs analysis relying on common EV markers (i.e. CD9, CD63 and CD81), however, a recent mass spectrometric study have shown the common EV markers were not as representative as one would expect of the total EV population in multiple cancers derived EVs(36). With the aid of single particle nanoflow analyzer and silica sizing beads, we could determine the size of B cells EVs which is very comparable to the TEM data, as well as the estimation of EVs secretion post activation that is in parallel to previous publications in a more robust and efficient manner(7, 22). These findings could be further applied in other types of immune EVs studies such as myeloid and lymphoid responses in research and clinical sectors.

One of the hallmarks of EVs are the expression of parental cell markers on their surface which allows EVs to be a novel immune responses progression monitoring tool(6). With the aid of the nanoflow analyzer, we determined the expression profile of EVs enriched tetraspanins marker, B cell marker, and CSR associated markers on B cell EVs during CSR. Using HEK293A EVs as internal negative control, we can confirm the antibodies specificity from the low background signals observed. Conventional tetraspanin EVs marker CD9 were expressed on 10% and 15% of total EVs released by stimulated CH12F3 and primary B cells respectively, suggesting studies relying on CD9 or other tetraspanins capture might not be conclusive in the total EVs analysis. CSR associated markers, IgG1 (primary B cells) and IgA (CH12F3) were enriched in EVs derived from switched parental cells at 96 hours post stimulation. Interestingly, IgM^+^IgG1^+^ and IgM^+^IgA^+^ EVs were detected in unswitched primary B cells and CH12F3 respectively at comparable levels (around 1 to 2% of total EVs; ∼10^6^ EVs) to switched parental cells, indicating that in the absence of stimulation, B cells undergo stochastic CSR without expressing switched isotypes antibodies (IgG1 or IgA) on their surface but incorporated on released EVs by some means. These findings highlight the biological relevance of B cell derived EVs during physiological CSR and also the potential of being further utilized as a novel biomarker for early detection of B cell associated autoimmune diseases such as hyper IgM syndrome and cancers.

EVs molecular cargo compositions have drawn much attention recently due to their ability to modulate intercellular signaling via targeted delivery of specific molecules such as RNA, DNA and proteins, particularly in ncRNA mediated gene regulation(6). In this study, ncRNA-seq was performed to understand the differential transcriptomic profile of B cell EVs during CSR as well as the variance of gene expression between parental cells and released EVs in both unswitched and switched states. From the complete DEG analysis and ImmGen database, we identified a novel CSR regulator, miR-5099, an uncharacterized miRNA that suppresses CSR via EVs intercellular signaling utilized by primary B cells, but not CH12F3. This data is indicating that healthy B cells suppress each other from hyperactivation via the delivery of EVs packaged miR-5099 between themselves, while the cancerous CH12F3 abolishes such machinery to increase hyperactivation. Gm26917, an uncharacterized lncRNA that was upregulated in CH12F3 EVs post stimulation, but not primary B cell EVs, it appeared to be a vital mediator for high CSR efficiency based on the Gm26917 silencing assay. Interestingly, Gm26917 is annotated as the top interacting target of miR-5099 from the ENCORI database, suggesting the high switching efficiency of CH12F3 possibly due to the high expression of Gm26917 and its suppression effect on miR-5099.

The mode of EVs mediated intercellular signaling could be very different depending on cell types, such as direct cargo transfer or receptor mediated signaling(37). Understanding the mode of action in each study model is vital for future studies’ direction. Interestingly, our data suggests EVs mediated CSR regulation in B cells are dependent on both EVs surface receptors and intraluminal ncRNAs cargo to different extent as seen in the loss of CSR regulatory phenotype in both proteinase-K and triton-X treated EVs compared to fresh EVs. Since B cells are professional antigen presenting cells within the immune system and producers of highly specific antibodies to neutralize pathogens or trigger professional effector cells(4), they are likely to benefit from these capacities to perform such strictly regulated signaling in autologous activation for CSR. Future studies and clinical applications could consider using B cells as primary EVs producers, given the possibilities of shuttling specific cargo of interest and engineering the surface antigen repertoire, to provide an unprecedented level of targeted drug delivery.

Following the identification of miR-5099 and Gm26917, the mRNA-seq analysis of anti-miR-5099 treated primary B cells revealed the connection of miR-5099 to ROD1/PTBP3, a RNA-binding protein that has been shown as an essential factor for AID binding to the switch locus for CSR to occur(28). PTBP gene family has been reported in multiple literatures in the context of B cell development and maintenance(29–31). Furthermore, our findings bolstered the indispensable role of ROD1 in B cell CSR and divulged a novel intercellular epigenomic signaling network associated to ROD1 that ultimately leads to CSR regulation. In the translational diagnostics perspective, the level of EVs miR-5099 circulating in the bloodstream can serve as a biomarker for early detection of B cells hyperactivation associated autoimmune diseases and blood cancers. Therapeutically speaking, with the ability to specifically package and isolate miR-5099 enriched EVs from B cells, EVs delivery of miR-5099 can ultimately be a form of drug treatment against hyperactivation of B cells associated autoimmune diseases and blood cancers.

Although our proposed novel EVs mediated CSR regulatory mechanism has provided a solid foundation and proof of concept for future studies in this direction, we believe there are far more unknown targets and alternative pathways underneath the full picture behind such EVs mediated epigenomic signaling in B cell CSR regulation but also other immune cells activation or suppression.

## AUTHORSHIP AND CONFLICT-OF-INTEREST STATEMENTS

The authors declare that they have no competing interests.

## ACKNOWLEDGEMENTS

We thank the Exeter Sequencing facility particularly Paul O’Neil and Karen Moore for help and advice. Christian Hacker and UZH microscopy facility for TEM support and advice. Bogdan Mateescu for discussions. Profs Matthew Scharff and Christian Münz for critical reading of the manuscript. RC is supported by SNF (310030_212553; 320030E_215576, CRSK-3_190550; IZSEZ0_204655; IZSEZ0_218166), Novartis Foundation (22B140), Vontobel Stiftung (41309), UZH-STWF (F-41309-01-01), and the UZH-URPP (Translational Cancer Research). KY is supported by a BioMedTech Entrepreneur Fellowship, BRIDGE-SNF (40B1-0_221565), and GRS-Innobooster (GRS-069/23).

## AUTHOR CONTRIBUTIONS

RC conceived the study. KY/RC designed the experiments. KY performed most experiments with some help from AA. KY and AA analyzed the data. KY and RC wrote the manuscript. All authors edited the manuscript. RC secured funding.

## DATA AND MATERIALS AVAILABILITY

Data is deposited in the GEO database.

## Notes

### Competing Interest Statement

The authors have declared no competing interest.

