## Supplementary material for "Intercellular epigenomic signaling during B cell maturation": Supp info

### SUPPLEMENTARY INFORMATION

| geneID | geneName | geneType | pairGeneID | pairGeneName | pairGeneType | interactionNum | expNum | seqTypeNum | totalReadsNum | FreeEnergy | AlignScore(Smith-Waterman) |
|---|---|---|---|---|---|---|---|---|---|---|---|
| MIMAT0020606 | mmu-miR-5099 | miRNA | ENSMUSG00000005566 | Trim28 | protein_coding | 1 | 1 | 1 | 2 | -34.8 | 35.5 |
| MIMAT0020606 | mmu-miR-5099 | miRNA | ENSMUSG00000006930 | Hap1 | protein_coding | 1 | 1 | 1 | 9 | -52.9 | 35 |
| MIMAT0020606 | mmu-miR-5099 | miRNA | ENSMUSG00000015656 | Hspa8 | protein_coding | 4 | 1 | 1 | 9 | -54.7 | 30 |
| MIMAT0020606 | mmu-miR-5099 | miRNA | ENSMUSG00000020571 | Pdia6 | protein_coding | 1 | 1 | 1 | 4 | -16.4 | 17 |
| MIMAT0020606 | mmu-miR-5099 | miRNA | ENSMUSG00000020812 | 1810032O08Rik | protein_coding | 1 | 1 | 1 | 4 | -34.1 | 32 |
| MIMAT0020606 | mmu-miR-5099 | miRNA | ENSMUSG00000022353 | Mtss1 | protein_coding | 1 | 1 | 1 | 3 | -57.5 | 37 |
| MIMAT0020606 | mmu-miR-5099 | miRNA | ENSMUSG00000024754 | Tmem2 | protein_coding | 1 | 1 | 1 | 2 | -37.1 | 21.5 |
| MIMAT0020606 | mmu-miR-5099 | miRNA | ENSMUSG00000028974 | Dffa | protein_coding | 1 | 1 | 1 | 1 | -36.5 | 18 |
| MIMAT0020606 | mmu-miR-5099 | miRNA | ENSMUSG00000033307 | Mif | protein_coding | 1 | 1 | 1 | 1 | -30.4 | 25.5 |
| MIMAT0020606 | mmu-miR-5099 | miRNA | ENSMUSG00000036435 | Exoc1 | protein_coding | 1 | 1 | 1 | 3 | -55.1 | 34.5 |
| MIMAT0020606 | mmu-miR-5099 | miRNA | ENSMUSG00000038482 | Tfdp1 | protein_coding | 1 | 1 | 1 | 2 | -43.9 | 26 |
| MIMAT0020606 | mmu-miR-5099 | miRNA | ENSMUSG00000044533 | Rps2 | protein_coding | 34 | 3 | 1 | 57 | -68.3 | 32.5 |
| MIMAT0020606 | mmu-miR-5099 | miRNA | ENSMUSG00000047030 | Spata2 | protein_coding | 1 | 1 | 1 | 2 | -48.6 | 32.5 |
| MIMAT0020606 | mmu-miR-5099 | miRNA | ENSMUSG00000050936 | Gm42743 | lincRNA | 1 | 1 | 1 | 3 | -42.5 | 23 |
| MIMAT0020606 | mmu-miR-5099 | miRNA | ENSMUSG00000055932 | Fto | protein_coding | 1 | 1 | 1 | 1 | -38.1 | 25.5 |
| MIMAT0020606 | mmu-miR-5099 | miRNA | ENSMUSG00000059995 | Atxn7l3 | protein_coding | 1 | 1 | 1 | 1 | -27.3 | 20 |
| MIMAT0020606 | mmu-miR-5099 | miRNA | ENSMUSG00000061724 | Gm2423 | processed_pseudogene | 1 | 1 | 1 | 1 | -37 | 23.5 |
| MIMAT0020606 | mmu-miR-5099 | miRNA | ENSMUSG00000064400 | Gm23927 | snoRNA | 2 | 1 | 1 | 4 | -67 | 31 |
| MIMAT0020606 | mmu-miR-5099 | miRNA | ENSMUSG00000064741 | Snord14a | snoRNA | 1 | 1 | 1 | 1 | -42.7 | 21.5 |
| MIMAT0020606 | mmu-miR-5099 | miRNA | ENSMUSG00000064776 | Gm25203 | snoRNA | 68 | 6 | 2 | 240 | -70.2 | 50.5 |
| MIMAT0020606 | mmu-miR-5099 | miRNA | ENSMUSG00000064791 | Snord14e | snoRNA | 1 | 1 | 1 | 2 | -43.8 | 28 |
| MIMAT0020606 | mmu-miR-5099 | miRNA | ENSMUSG00000064923 | Gm22042 | snRNA | 1 | 1 | 1 | 2 | -50.1 | 28 |
| MIMAT0020606 | mmu-miR-5099 | miRNA | ENSMUSG00000065094 | Snord1a | snoRNA | 1 | 1 | 1 | 4 | -34.1 | 32 |
| MIMAT0020606 | mmu-miR-5099 | miRNA | ENSMUSG00000065176 | Rnu12 | snRNA | 1 | 1 | 1 | 1 | -37.1 | 15.5 |
| MIMAT0020606 | mmu-miR-5099 | miRNA | ENSMUSG00000065304 | Gm23245 | snoRNA | 1 | 1 | 1 | 1 | -31.3 | 25.5 |
| MIMAT0020606 | mmu-miR-5099 | miRNA | ENSMUSG00000065870 | Rnu3a | snoRNA | 22 | 4 | 1 | 37 | -64.1 | 35.5 |
| MIMAT0020606 | mmu-miR-5099 | miRNA | ENSMUSG00000065883 | Gm22865 | snoRNA | 11 | 2 | 1 | 37 | -64.6 | 30.5 |
| MIMAT0020606 | mmu-miR-5099 | miRNA | ENSMUSG00000073867 | AA474408 | TEC | 1 | 1 | 1 | 1 | -33.4 | 24 |
| MIMAT0020606 | mmu-miR-5099 | miRNA | ENSMUSG00000077709 | Snora64 | snoRNA | 34 | 3 | 1 | 57 | -68.3 | 32.5 |
| MIMAT0020606 | mmu-miR-5099 | miRNA | ENSMUSG00000080610 | Snord110 | snoRNA | 1 | 1 | 1 | 1 | -40.6 | 32.5 |
| MIMAT0020606 | mmu-miR-5099 | miRNA | ENSMUSG00000080615 | Snord99 | snoRNA | 1 | 1 | 1 | 2 | -39.9 | 19.5 |
| MIMAT0020606 | mmu-miR-5099 | miRNA | ENSMUSG00000083563 | Gm13340 | unprocessed_pseudogene | 1 | 1 | 1 | 3 | -28.5 | 24 |
| MIMAT0020606 | mmu-miR-5099 | miRNA | ENSMUSG00000086290 | Snhg12 | lincRNA | 1 | 1 | 1 | 2 | -39.9 | 19.5 |
| MIMAT0020606 | mmu-miR-5099 | miRNA | ENSMUSG00000087943 | Gm24245 | miRNA | 7 | 2 | 1 | 12 | -66.7 | 33.5 |
| MIMAT0020606 | mmu-miR-5099 | miRNA | ENSMUSG00000088088 | Rmrp | ribozyme | 1 | 1 | 1 | 3 | -40.7 | 30.5 |
| MIMAT0020606 | mmu-miR-5099 | miRNA | ENSMUSG00000088252 | Snord13 | snoRNA | 1 | 1 | 1 | 1 | -30.9 | 23.5 |
| MIMAT0020606 | mmu-miR-5099 | miRNA | ENSMUSG00000088895 | Gm22519 | snoRNA | 1 | 1 | 1 | 1 | -31.6 | 17.5 |
| MIMAT0020606 | mmu-miR-5099 | miRNA | ENSMUSG00000089417 | Gm22009 | scaRNA | 1 | 1 | 1 | 2 | -27.9 | 20 |
| MIMAT0020606 | mmu-miR-5099 | miRNA | ENSMUSG00000090862 | Rps13 | protein_coding | 1 | 1 | 1 | 1 | -42.7 | 21.5 |
| MIMAT0020606 | mmu-miR-5099 | miRNA | ENSMUSG00000092674 | Gm24105 | misc_RNA | 1 | 1 | 1 | 3 | -40.5 | 22 |
| MIMAT0020606 | mmu-miR-5099 | miRNA | ENSMUSG00000092730 | Snora24 | snoRNA | 1 | 1 | 1 | 1 | -51.5 | 44 |
| MIMAT0020606 | mmu-miR-5099 | miRNA | ENSMUSG00000092909 | Gm25732 | miRNA | 1 | 1 | 1 | 1 | -50.6 | 35.5 |
| MIMAT0020606 | mmu-miR-5099 | miRNA | ENSMUSG00000094405 | Gm23143 | snRNA | 1 | 1 | 1 | 2 | -41.4 | 23 |
| MIMAT0020606 | mmu-miR-5099 | miRNA | ENSMUSG00000094655 | Gm25360 | snRNA | 1 | 1 | 1 | 1 | -36.8 | 25 |
| MIMAT0020606 | mmu-miR-5099 | miRNA | ENSMUSG00000094916 | Gm26391 | rRNA | 1 | 1 | 1 | 1 | -47.5 | 26 |
| MIMAT0020606 | mmu-miR-5099 | miRNA | ENSMUSG00000095118 | Snord14d | snoRNA | 3 | 1 | 1 | 7 | -54.7 | 30 |
| MIMAT0020606 | mmu-miR-5099 | miRNA | ENSMUSG00000095217 | Hist1h2bn | protein_coding | 1 | 1 | 1 | 2 | -48.3 | 27 |
| MIMAT0020606 | mmu-miR-5099 | miRNA | ENSMUSG00000095590 | Gm24305 | snRNA | 1 | 1 | 1 | 1 | -44.9 | 26 |
| MIMAT0020606 | mmu-miR-5099 | miRNA | ENSMUSG00000095738 | Gm25313 | snRNA | 1 | 1 | 1 | 1 | -24.7 | 20 |
| MIMAT0020606 | mmu-miR-5099 | miRNA | ENSMUSG00000098425 | Gm24949 | snoRNA | 42 | 2 | 1 | 85 | -72.1 | 36 |
| MIMAT0020606 | mmu-miR-5099 | miRNA | ENSMUSG00000098943 | Rnu3b1 | snoRNA | 20 | 2 | 1 | 27 | -73.2 | 33.5 |
| MIMAT0020606 | mmu-miR-5099 | miRNA | ENSMUSG00000099291 | Rnu3b3 | snoRNA | 19 | 1 | 1 | 30 | -73.8 | 40.5 |
| MIMAT0020606 | mmu-miR-5099 | miRNA | ENSMUSG00000104856 | Rnu3b3 | snoRNA | 19 | 1 | 1 | 30 | -73.8 | 40.5 |
| MIMAT0020606 | mmu-miR-5099 | miRNA | ENSMUSG00000104921 | Gm23927 | snoRNA | 2 | 1 | 1 | 4 | -67 | 31 |
| MIMAT0020606 | mmu-miR-5099 | miRNA | ENSMUSG00000104960 | Snhg8 | processed_transcript | 1 | 1 | 1 | 1 | -51.5 | 44 |
| MIMAT0020606 | mmu-miR-5099 | miRNA | ENSMUSG00000105025 | Rnu3b1 | snoRNA | 20 | 2 | 1 | 27 | -73.2 | 33.5 |
| MIMAT0020606 | mmu-miR-5099 | miRNA | ENSMUSG00000105037 | Gm25203 | snoRNA | 68 | 6 | 2 | 240 | -70.2 | 50.5 |
| MIMAT0020606 | mmu-miR-5099 | miRNA | ENSMUSG00000105790 | AC122546.1 | miRNA | 1 | 1 | 1 | 3 | -40.5 | 22 |
| MIMAT0020606 | mmu-miR-5099 | miRNA | ENSMUSG00000105827 | Hist2h2bb | protein_coding | 1 | 1 | 1 | 3 | -42.5 | 23 |
| MIMAT0020606 | mmu-miR-5099 | miRNA | ENSMUSG00000106147 | Rnu3a | snoRNA | 22 | 4 | 1 | 37 | -64.1 | 35.5 |
| MIMAT0020606 | mmu-miR-5099 | miRNA | ENSMUSG00000115420 | AL732506.1 | lincRNA | 1 | 1 | 1 | 3 | -40.7 | 30.5 |
| MIMAT0020606 | mmu-miR-5099 | miRNA | ENSMUSG00000018707 | Dync1h1 | protein_coding | 1 | 1 | 1 | 4 | -50.7 | 31 |
| MIMAT0020606 | mmu-miR-5099 | miRNA | ENSMUSG00000064570 | Gm25800 | snoRNA | 1 | 1 | 1 | 1 | -43.9 | 24 |
| MIMAT0020606 | mmu-miR-5099 | miRNA | ENSMUSG00000065715 | Snord7 | snoRNA | 1 | 1 | 1 | 1 | -48.1 | 38 |
| MIMAT0020606 | mmu-miR-5099 | miRNA | ENSMUSG00000076258 | Gm23935 | miRNA | 1 | 1 | 1 | 1 | -98.8 | 33.5 |
| MIMAT0020606 | mmu-miR-5099 | miRNA | ENSMUSG00000097971 | Gm26917 | lincRNA | 70 | 1 | 1 | 108 | -70.1 | 42 |
| MIMAT0020606 | mmu-miR-5099 | miRNA | ENSMUSG00000098178 | Gm42418 | lincRNA | 67 | 4 | 1 | 96 | -87.3 | 37.5 |
| MIMAT0020606 | mmu-miR-5099 | miRNA | ENSMUSG00000098973 | Mir6236 | miRNA | 1 | 1 | 1 | 1 | -37.2 | 24 |
| MIMAT0020606 | mmu-miR-5099 | miRNA | ENSMUSG00000114081 | Gm48904 | TEC | 1 | 1 | 1 | 1 | -37.7 | 19.5 |
| MIMAT0020606 | mmu-miR-5099 | miRNA | MIMAT0024857 | mmu-miR-6236 | miRNA | 1 | 1 | 1 | 1 | -37.2 | 24 |

**Table S1**. List of RNA interactive targets of miR-5099 .

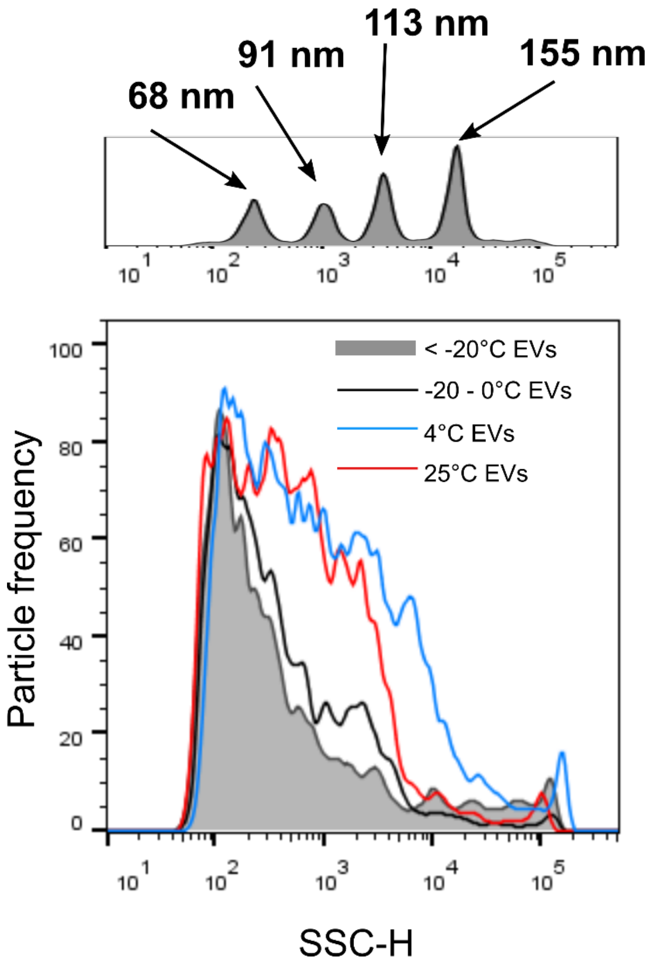


**Figure S1**. Depletion of intact EVs via multiple cycles of freeze-thaw cycles. Representative side scatter dot plot of CH12F3 EVs subjected at least 3 cycles of freezing at -80°C (solid grey), -20°C (grey line) and positive control of EVs stored at 4°C (blue), and 25°C (red).

S1

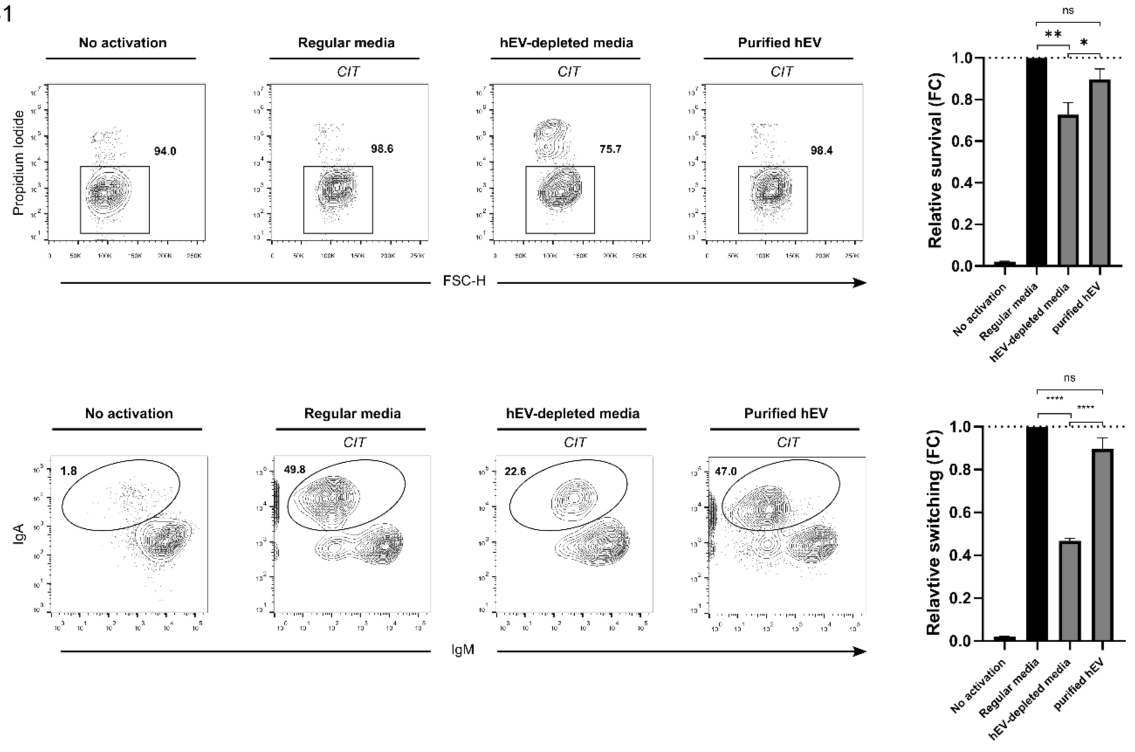


**Figure S2**. Fetal calf serum derived heterologous EVs affect B cell survival and CSR efficiency. Representative FACS plots and quantifications of viability and CSR efficiency in CH12F3 in regular media, FCS hEVs depleted media, addition of purified FCS hEVs in depleted media (left to right). Representative of three experiments, mean ±SD shown in columns by one-way ANOVA with Tukey's multiple comparison test (*p<0.05; **p < 0.01; ***p < 0.001).

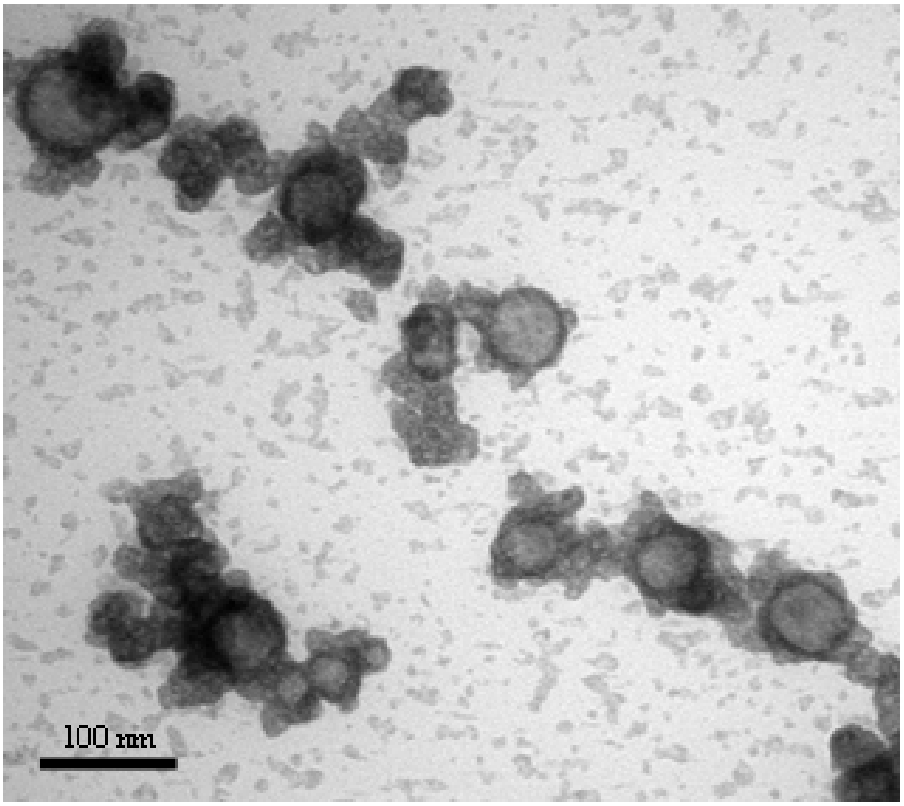


**Figure S3**. Transmission electron microscopy of purified EVs derived from CH12F3.

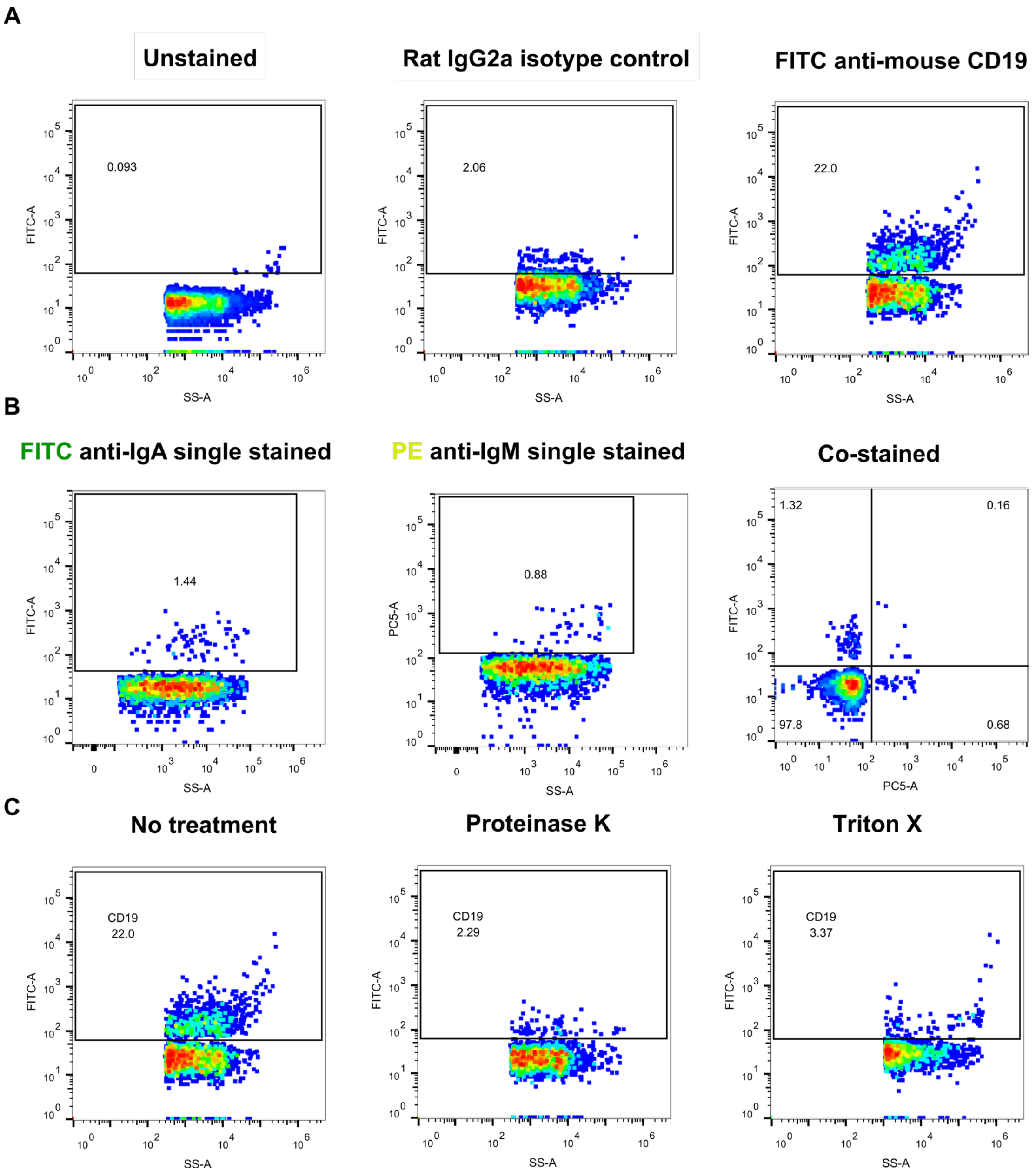


**Figure S4**. Representative nanoflow dot plot of CH12F3 EVs (unstained, isotype control, FITC-anti-mouse CD19). (A) Antibodies isotype control for EVs nanoflow analysis. (B) Single-stained controls for double-stained (FITC & PE) EVs nanoflow analysis. (C) Quality assessment of proteinase-K and 1% triton-X treatment of CH12F3 EVs prior to functional assays.

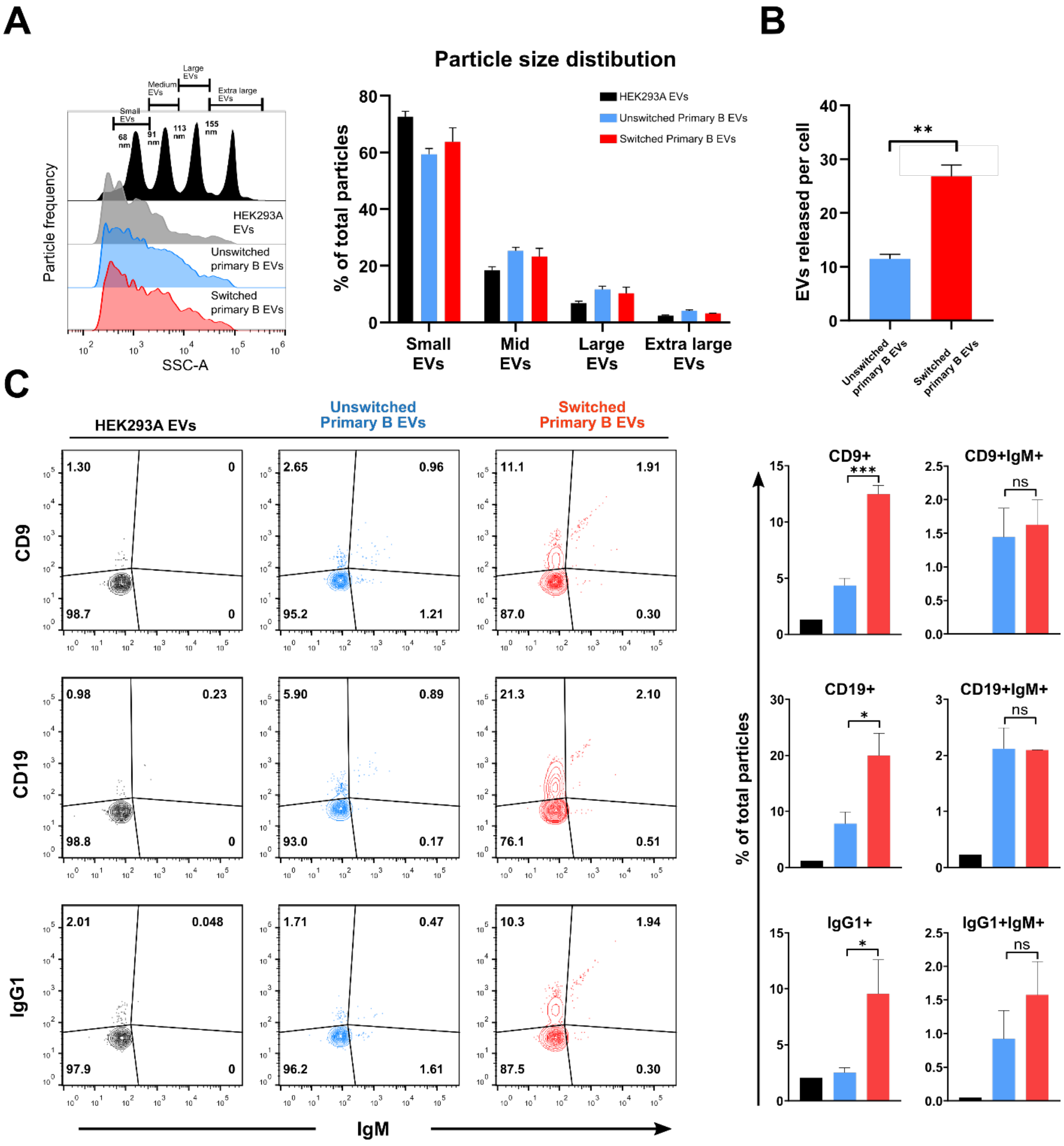


**Figure S5**. Multiplex characterization of primary B cell EVs during CSR.

(A) Representative side scatter plot and estimated size distribution quantification of primary B EVs during CSR, using silica sizing beads with mix of 66 nm (small), 91 nm (medium), 113 nm (large), and 155 nm (extra-large) (r index = 1.46). (B) Quantification of estimated EVs particles released per cell via dividing the total number of particles by number of parental cells. (C) Representative FACS plots and quantification of denoted markers of primary B EVs during CSR. Representative of three experiments, mean ±SD shown in columns by one-way ANOVA with Tukey's multiple comparison test (*p<0.05; **p < 0.01; ***p < 0.001).

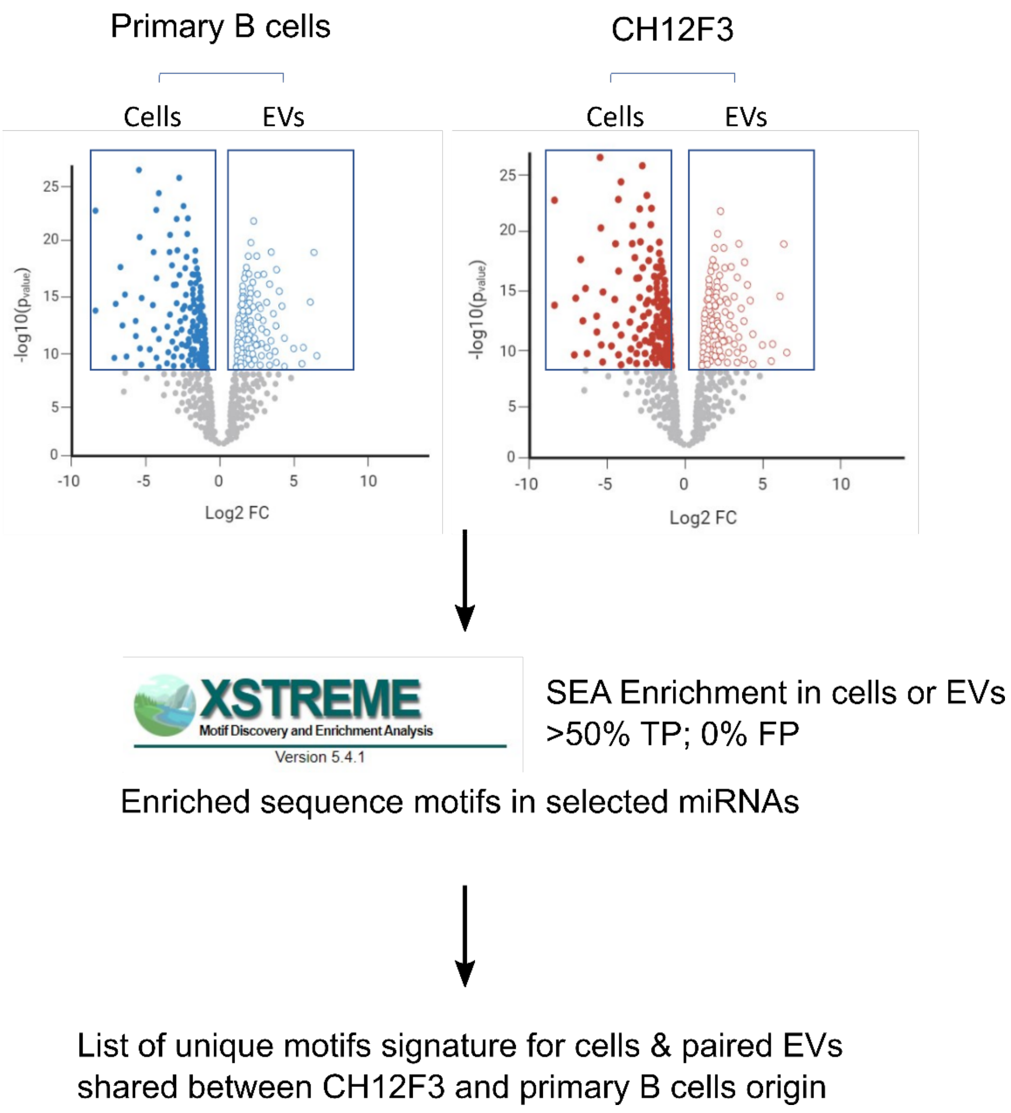


**Figure S6.** Schematic outline of motif enrichment analysis.

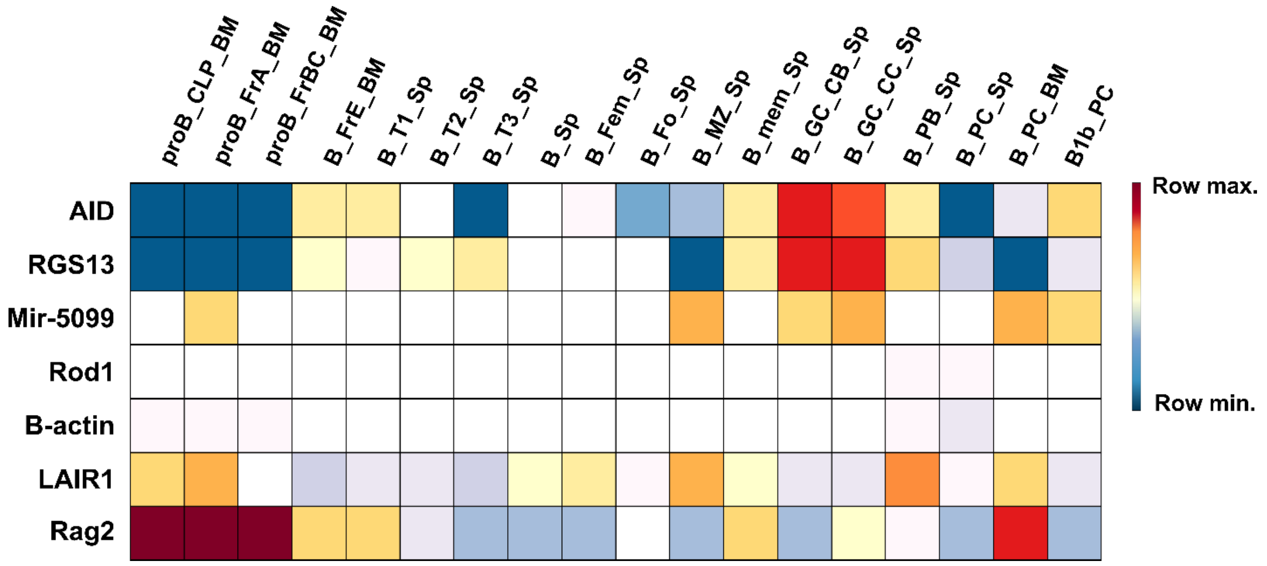


**Figure S7**. In vivo expression of miR-5099 in various germinal center (GC) B cell compartment. Heatmap showing the in vivo expression of denoted genes in various B cell compartments. Positive control for GC expression: AID and TGS13; Housekeeping gene: B actin; Negative control for GC expression: LAIR1 and RAG2.

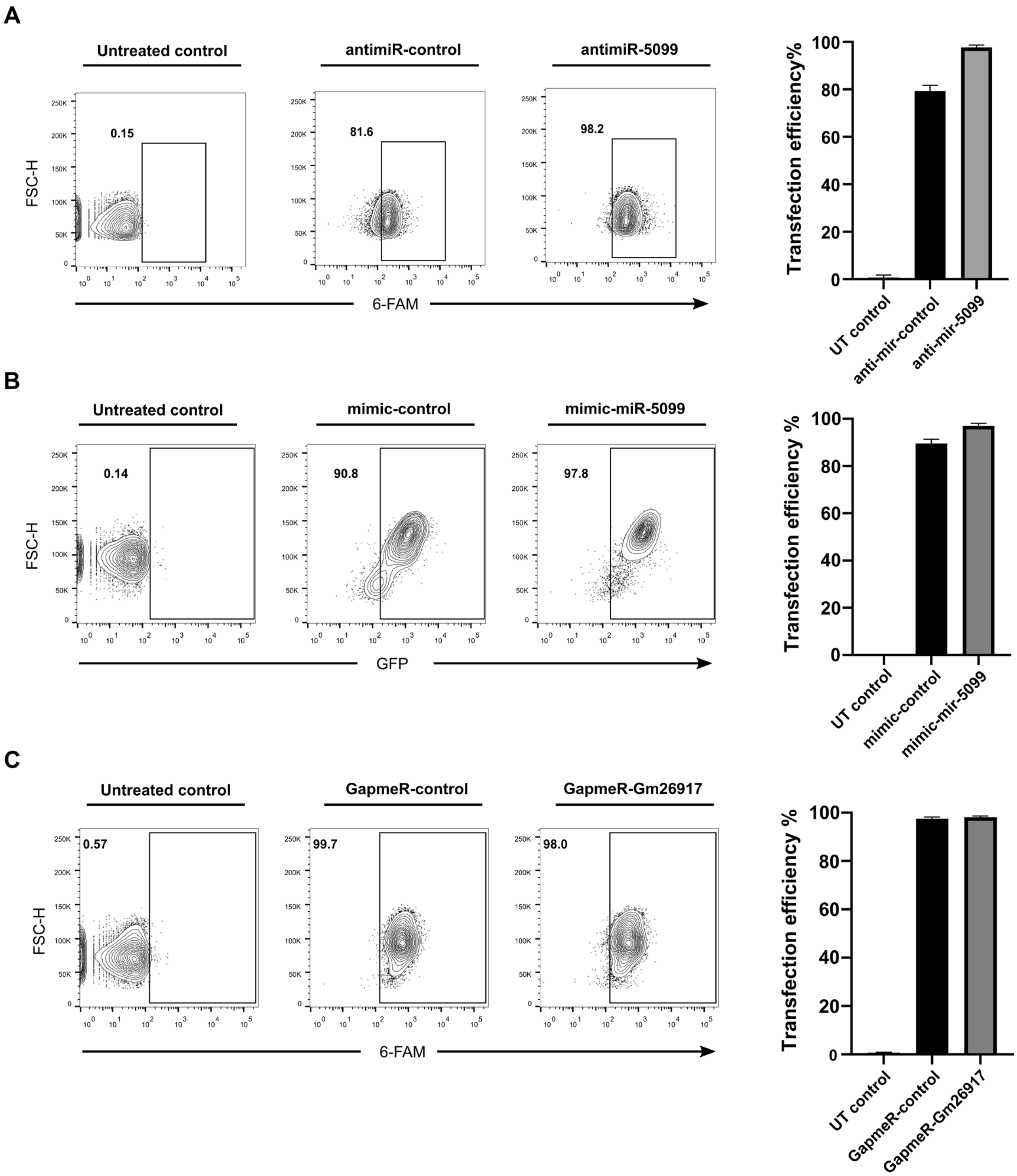


**Figure S8**. Antagomir/GapmeR direct uptake and mimic transfection efficiency assessment in CH12F3. Representative FACS plot and quantification of 6-Carboxyfluorescein (6-FAM)/GFP signals in CH12F3 with (A) 6-FAM conjugated anti-mir-control or anti-miR-5099, (B) GFP plasmid co-transfected with mimic-control or mimic-mir-5099, and (C) 6-FAM conjugated GapmeR-control or GapmeR-Gm26917. Representative of three experiments, mean ±SD shown in columns by one-way ANOVA with Tukey's multiple comparison test (*p<0.05; **p < 0.01; ***p < 0.001).

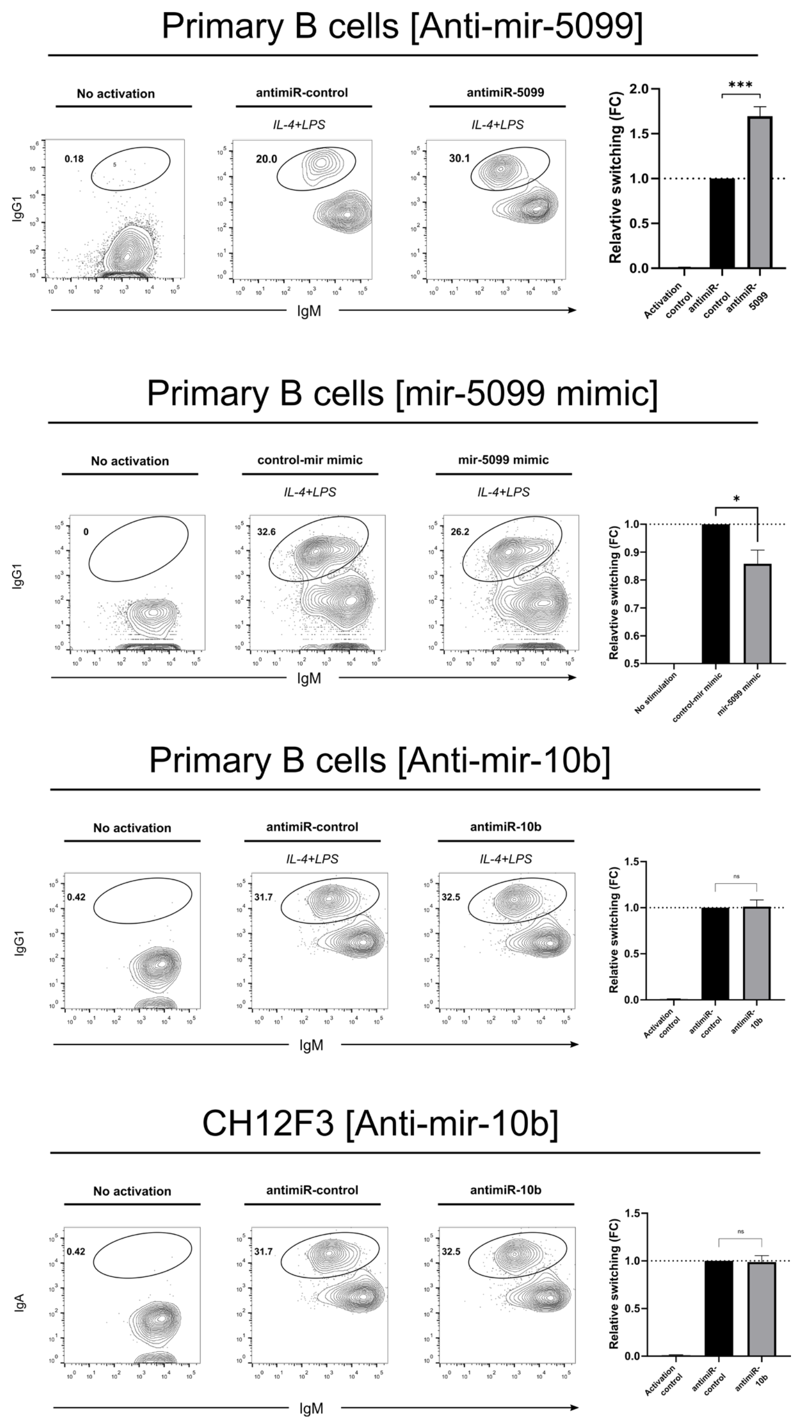


**Figure S9**. Representative FACS plots and quantifications of CSR efficiency in primary B cells (top 2) treated with anti-miR-5099, miR-5099 mimic and primary B cells and CH12F3 (bottom 2) treated with anti-mir-10b-5p compared to respective negative controls. Representative of three experiments, mean ±SD shown in columns by one-way ANOVA with Tukey's multiple comparison test (*p<0.05; **p < 0.01; ***p < 0.001).

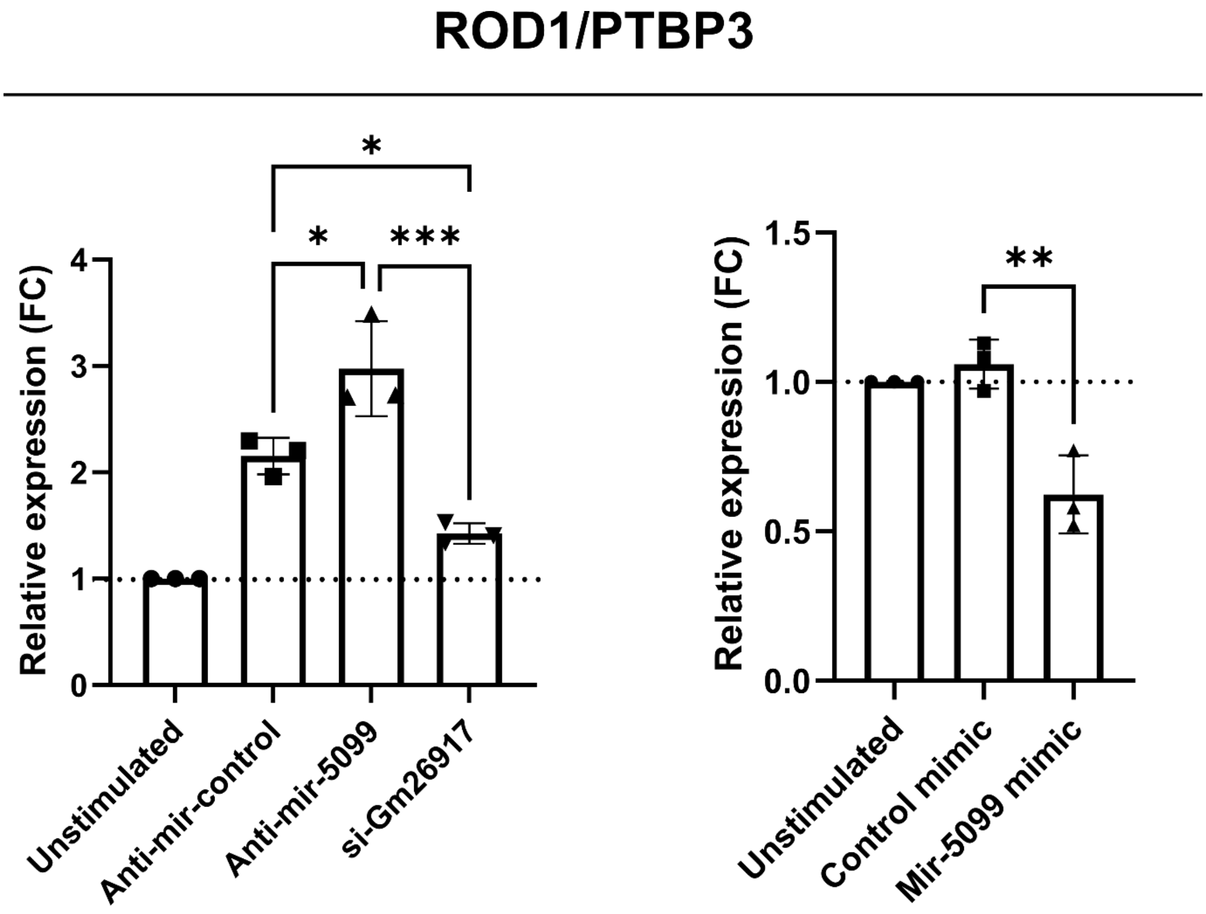


**Figure S10**. Quantifications of relative expression of ROD1/PTBP3 in CH12F3 treated with anti-miR-5099, miR-5099 mimic, and GapmeR Gm26917 compared to control. Mean ±SD shown in columns by one-way ANOVA with Tukey’s multiple comparison test (*p<0.05; **p < 0.01; ***p < 0.001).

| Primer | Sequence |
|---|---|
| mRod1-F | CTCGCTTAGACCTTCCTACTGG |
| mRod1-R | CTGCTTGAGGAAATGCGATGGC |
| mAID-F | GAAAGTCACGCTGGAGACCG |
| mAID-R | TCTCATGCCGTCCCTTGG |
| mSwitch alpha-F | GACATGATCACAGGCACAGG |
| mSwitch alpha-R | TTCCCCAGGTCACATTCATCGT |
| mouse gamma1-GLT-F | CGAGAAGCCTGAGGAATGTGT |
| mouse gamma1-GLT-R | GGAGTTAGTTTGGGCAGCAGAT |
| mouse mu-GLT-F | TCTGGACCTCTCCGAAACCA |
| mouse mu-GLT-R | ATGGCCACCAGATTCTTATCAGA |
| mouse actb-F | CGTGAAAAGATGACCCAGATCA |
| mouse actb-R | TGGTACGACCAGAGGCATACAG |

**Table S2.** Primers sequence for RT-qPCR.
